# Leaf traits in *P*h*aseolus vulgaris* and *Phaseolus acutifolius* reveal divergent terminal drought coping strategies

**DOI:** 10.1101/2025.10.28.685217

**Authors:** A. Violeta Salazar-Chavarría, Guadalupe T. Zavala-Padilla, Arely Lechuga-Jiménez, V. Miguel Palomar, Alexis Acosta-Maspons, Alejandra A. Covarrubias

**Author notes:** **Corresponding author:** Alejandra A. Covarrubias.

## Abstract

*Phaseolus* beans play a crucial role in global food security, providing a sustainable source of protein and micronutrients for diverse populations. However, their productivity is severely affected by prolonged drought during the reproductive stage (terminal drought), which drastically reduces grain yield and quality. While substantial progress has been made in understanding plant drought responses during the vegetative stage, less is known about the responses during the reproductive phase. Among the different responses, little is known about the accompanying anatomical and physiological changes across plant organs. In particular, the leaf plasticity in response to environmental fluctuations remains unexplored under terminal drought. This study investigates the anatomical, physiological and molecular leaf responses to terminal drought in two resistant genotypes of *Phaseolus vulgaris* (common bean) and *Phaseolus acutifolius* (tepary bean). Despite comparable water status under stress, tepary mature leaves present longer major veins, larger xylem area, increased air space, and thicker cuticles than common bean. These traits likely contribute to improve water transport and gas exchange. Consistently, tepary bean maintained higher photosynthetic performance under well-watered and drought conditions, with considerable superior carbon fixation rates under irrigation. Elevated starch and sucrose accumulation in tepary leaves under both treatments further supports its enhanced carbon assimilation. RNAseq analysis indicated that some of these traits are partly transcriptomic dependent. Together, our findings highlight the diverse leaf-level adaptations that underlie terminal drought resistance in these species. The enhance anatomical and physiological traits in tepary bean offer valuable insights for improving drought resilience in common bean in a changing climate.

**Highlight:** Leaf vein structure, xylem vessels density, and cuticle thickness enhance carbon assimilation and functional maintenance in tepary bean under drought during the reproductive stage, revealing key traits for common bean improvement.

## Introduction

In recent decades, climate change has intensified meteorological phenomena such as heat waves and prolonged droughts, causing severe economic losses (van Daalen *et al*., 2022; Vicente-Serrano *et al*., 2022). Water scarcity is a major limitation for crop growth and yield, triggering molecular, physiological, and anatomical responses in plants. Although molecular and physiological mechanisms have been extensively studied, anatomical changes remain less understood. Reported effects of water deficit include alterations in tissue thickness and size in roots, stems, and leaves, particularly within the vascular system (Mano *et al*., 2023; Saeed *et al*., 2016; Shafqat *et al*., 2021; Velikova *et al*., 2020; Vital *et al*., 2022). Leaves, in particular, display notable plasticity modifying thickness, area, and dry matter content, in association with changes in cell size and cell wall properties (Dovrat *et al*., 2019; Flexas *et al*., 2021; Funk *et al*., 2021; Sack and Scoffoni, 2013; Yavas *et al*., 2024). Alterations in airspace proportion have also been reported (de Avila Silva *et al*., 2021; Huang *et al*., 2019; Mendes *et al*., 2022), affecting fundamental physiological processes.

Photosynthesis is one of the physiological processes most affected by water deficit, primarily due to reduced water content and CO_2_ availability. The decrease in CO_2_ under drought conditions results from both stomatal and non-stomatal restraints (Gimeno *et al*., 2019; Grassi and Magnani, 2005; Varone *et al*., 2012). Stomatal limitations depend on stomatal characteristics and conductance, while non-stomatal factors involve mesophyll conductance and other physiological processes regulating CO_2_ transport to the chloroplast (Flexas and Medrano, 2002; Gago *et al*., 2019; Gimeno *et al*., 2019; Nadal and Flexas, 2018). These processes are strongly influenced by leaf anatomy, which determines the diffusion pathway of CO_2_ from the sub-stomatal cavity to the stroma. Critical anatomical traits include the proportion of air spaces, cell wall thickness, and the size and arrangement of the parenchyma cells (Clarke *et al*., 2021; Evans, 2021; Han *et al.*, 2018; Mizokami *et al*., 2019). Variations in these features can enhance CO_2_ delivery to fixation sites, particularly when stomatal closure limits gas exchange during drought (Cano *et al*., 2013; Han *et al*., 2019).

Water content in plant tissues is another essential factor for maintaining efficient photosynthetic activity. Its proper distribution largely depends on the vascular system. Duirng drought, some plant species exhibit modifications in the density and size of major and minor veins, as well as in the vessels that compose them, thus improving water transport and diffusion (Li *et al*., 2022). Such anatomical adjustments have been associated with drought tolerance in species like Acer and Quercus, which typically grow in environments with limited water availability (Nardini *et al*., 2012). Vessel size, in particular, represent an adaptive trait under water deficit, with contrasting responses observed among species (Haworth *et al*., 2017; Nogueira *et al*., 2020; Rodríguez-Ramírez *et al*., 2022).

Drought is one of the most recurrent and severe environmental stresses affecting plants. When it occurs during flowering and grain-filling stages, referred to as terminal drought, it can drastically reduce crop quality and yield (Acosta-Gallegos and Shibata, 1989; Farooq *et al*., 2014). The impact is particularly severe under rainfed conditions, as is common for bean cultivation in developing countries (Broughton *et al*., 2003), where terminal drought can cause yield losses from 70 to 100%, depending on the intenisty and duration of the dry season (Acosta-Gallegos and Shibata, 1989; Smith *et al*., 2019). Common bean (*Phaseolus vulgaris*) is among the most socio-economically important crops worldwide, both in terms of cultivated area and nutritional value, as it provides a rich source of protein, starch, unsaturated fatty acids, dietary fiber, vitamins, and minerals such as iron and zinc (Ganesan and Xu, 2017; Gepts *et al*., 2008). Consequently, several drought-resistant cultivars have been developed, including ‘Pinto Saltillo’, a member of the Durango race, which maintains seed production even under severe water deficit (González-Lemes *et al*., 2023). Another cultivated species capable of sustaining yield under drought conditions is *Phaseolus acutifolius* (tepary bean), a bean native to arid regions in northern Mexico and the southwestern United States. This species has a short life cycle and displays remarkable tolerance not only to drought but also to extreme temperatures, salinity, acidic soils, and nutrient deficiencies (Beebe *et al*., 2013; Micheletto *et al*., 2007; Moghaddam *et al.*, 2021; Suarez *et al*., 2022).

To understand traits associated to drought resistance in genotypes valuable for crop improvement, previous work demonstrated that under terminal drought treatment, the *P. vulgaris* cultivar ‘Pinto Saltillo’ closes early (isohydric strategy), while *P. acutifolius* tepary bean (TARS-Tep 32) stomatal opening longer (anisohydric strategy) (Polania *et al*., 2022). Tepary bean’s shoot biomass was less affected (38% vs. 42% reduction in ‘Pinto Saltillo’), and its photosynthetic performance was superior (Polania *et al*., 2022). To explore potential anatomical bases, we analyzed mesophyll organization, xylem area and vessels, major vein length and cuticle thickness. Tepary bean displayed longer major veins, larger xylem area, thicker cuticle, and greater intercellular air space in mature leaves. Despite similar leaf water status, TARS-Tep 32 achieved more efficient carbon assimilation than ‘Pinto Saltillo’ under both well-watered and terminal drought conditions. Comparative transcriptomic analysis revealed more differential abundant transcripts (DAT) in *P. vulgaris* leaves, suggesting that ‘Pinto Saltillo’ experiences higher stress than tepary bean. The cultivars of both species, however, shared up-regulated transcripts associated to water deficit responses, including those encoding LEA proteins, chaperones, antioxidant enzymes, and abscisic acid-responsive proteins. In *P. acutifolius* leaves, enriched up-regulated transcripts were primarily associated to cutin and wax biosynthesis, carbohydrate and lipid metabolism, cell wall remodeling, and development. In contrast, *P. vulgaris* showed stronger enrichment of stress-responsive transcripts, as well as transcripts involved in amino acid metabolism and in transport and plasma-membrane turnover. Several drought-responsive transcripts corresponded to the characteristic anatomical and physiological traits of species, linking molecular responses to functional drought adaptations.

Overall, our findings reveal diverse leaf-level adaptations and adjustments associated with terminal drought resistance in common and tepary beans, providing valuable insights for improving crop resilience under climate change.

## Materials and Methods

### Plant materials and growing conditions

All experiments were performed using two drought tolerant elite genotypes from *Phaseolus vulgaris* L. (common bean) cultivar ‘Pinto Saltillo’ (hereinafter referred as *Pv*), an elite genotype from Durango race (Rosales *et al*., 2013), and *Phaseolus acutifolius* A. Gray (Tepary bean), line TARS-Tep 32 (hereinafter referred as *Pa*), a cultivar improved for heat and terminal drought resistance (Barrera *et al*., 2024; Porch *et al*., 2013; Porch *et al.*, 2017). These genotypes were kindly provided by Dr. Jorge Acosta-Gallegos (Instituto Nacional de Investigaciones Agrícolas y Pecuarias, INIFAP, México) and by Dr. Tim Porch (United States Department of Agriculture, USDA, Puerto Rico), respectively. Seeds were sterilized, scarified and germinated in water saturated filter paper in Petri dishes. Selected germinated seeds were transferred to pots (21 cm X 16 cm X 15.5 cm, 5 L) containing vermiculite-coarse 3A/perlite/Metromix-Mix 3 Sunshine (4:3:3). We followed a randomized complete block experimental design with one plant per pot and at least seven biological replicates. After sowing and at the flowering onset, we added Osmocote Plus (15-9-12 NPK), an organic fertilizer. Plants were grown under greenhouse-controlled conditions with an average temperature of 25 ± 2 °C, a relative humidity of approximately 60% and a natural light/dark cycle at the Instituto de Biotecnología (IBt) of the Universidad Nacional Autónoma de México (UNAM) in Cuernavaca, Morelos, México, (18°59ʹ24″ N, 99°14ʹ3″ W) during 2022-2024. After sowing, irrigation was carefully performed, by daily watering to keep substrate at field capacity during plant vegetative development until flowering onset (approx. 30 days after sowing, depending on the genotype phenology). At this stage, plants were divided into two groups: one group, the well-watered (WW) set, where plants were maintained at field capacity, and a second group, where plants were subjected to a water deficit treatment by keeping the substrate at 25% of field capacity to simulate terminal drought (TD). This treatment was applied from flowering to harvest, which was performed at the mid-pod filling stage.

### Gas exchange measurements and photosynthetic parameters

All measurements were performed in completely expanded mature leaves from plants at the mid-pod filling stage. The mature leaves analyzed were those whose development and per treatment from at least six plants in each case. Using an infrared gas analyzer (LI-6400; LI-COR), we determined net photosynthesis rate (An) and intercellular CO_2_ concentration (Ci). We applied in a leaf chamber a saturating light intensity of 1000 µmol m^−2^ s^−1^, an air flow of 400 µmol s^−1^ and 400 µmol CO_2_. Leaf relative chlorophyll content (SPAD units) and the photosynthetic efficiency of photosystem II in light-adapted leaves (Fv/Fm ratio) were determined in fully expanded leaves using a MultispeQ (PhotosynQ; (Kuhlgert *et al*., 2016). All measurements were performed between 11 am and 1 pm.

### Determination of CO_2_ response curves (A_n_/C_i_) and maximum carboxylation rate (V_cmax_)

To determine the photosynthetic CO_2_ responses, the net CO_2_ assimilation rate (A_n_) was measured in response to different CO_2_ concentrations. For this, three mature leaves from each species were collected under both experimental conditions. Measurements were conducted between 10 am and 12 pm using an infrared gas analyzer (LI-6400. LI-COR, Inc., Lincoln, NE USA). CO_2_ response curves were obtained using CO_2_ concentrations of 50,100, 200, 400, 600, 800 y 1000 µmol of CO_2_, with an air flow of 400 µmol s^−1^ and a saturating light intensity of 1000 µmol m^−2^ s^−1^. The A_n_/C_i_ curves were fitted using the bilinear method implemented in the ‘fitaci’ function of the *plantecophys* R package, which models leaf gas exhange data (Duursma, 2015). This approach is based on the Farquhar-Berry-von Caemmerer-Berry (FvCB) model of photosynthesis generally described as the following equation: A_n_ = *min* (A_c_, A_j_) – Rd, where A_n_ is the net CO_2_ assimilation rate, A_c_ and A_j_ are the gross photosynthesis rates under limiting Rubisco activity or limiting Rubisco regeneration, respectively, and Rd is the rate of non-photorespiratory mitochondrial respiration in the light. The term *min* means that the A_n_ is determined by the minimum of the two possible limiting processes A_c_ or A_j_ (Farquhar *et al*., 1980).

The R package *plantecophys* estimates the photosynthesis rate under limiting Rubisco activity (*Ac*) using the following relation:

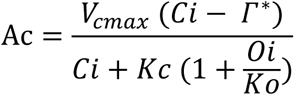

Where *V_cmax_* is the maximum rate of Rubisco activity; *Ci* and *Oi* are the intercellular concentrations of CO_2_ and O_2_, respectively; *Kc* and *Ko* are the Michaelis-Menten constants of Rubisco for CO_2_ and O_2_, respectively; and ***Γ***\* is the CO_2_ compensation point in the absence of mitochondrial respiration (Duursma, 2015; Egesa *et al*., 2024).

For the calculation of photosynthesis rate under limiting Rubisco regeneration activity (*Aj*) this package uses the following equation:

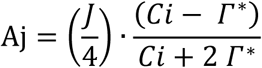

Where *J* is the rate of electron transport, *Ci* is the intercellular CO_2_ concentration and ****Γ***\** is the CO_2_ compensation point in the absence of mitochondrial respiration (Duursma, 2015; Song *et al*., 2021).

For *V_cmax_* calculation, the *plantecophys* R package only considers the initial portion of the A_n_/C_i_ curve (C_i_ < 400 μmol CO_2_), normalizing the data with the leaf temperature (25 °C), using the Arrhenius equation as follows:

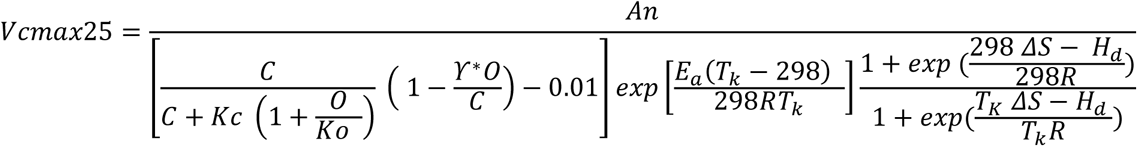

Where C is the CO_2_ concentration in the bundle sheath; *K_c_* and *K_o_* are the Michaelis-Menten constants of Rubisco for CO_2_ and O_2_, respectively; *O* is the partial pressure of O_2_; *γ\** is half the reciprocal of Rubisco specificity; *E_a_* is the activation energy; *T_k_* is the leaf temperature in Kelvin degrees; *R* is the universal gas constant; *ΔS* is the entropy term; and *H_d_* is the deactivation energy (Song *et al*., 2021).

### Determination of fructose, glucose, sucrose, and starch content

Three mature leaves from each species were collected under both experimental conditions. The samples were immediately frozen and ground in liquid nitrogen to a fine powder. For each extraction, 80-100 mg of tissue were used. Starch quantification was carried out in two phases. First, soluble sugars were extracted following the method described by Gonzalez-Lemes *et al*. (González-Lemes *et al*., 2023). After sugar removal, the remaining dry pellet was resuspended in 500 μL of amylase buffer (50 mM Tris-HCl, 2 mM CaCl_2_, 150 mM NaCl, pH 7.0) and incubated in a Thermomix Compact (Eppendorf) at 400 rpm and 95 °C for 90 min to initiate starch hydrolysis. Samples were then centrifuged at 15,000 rpm for 10 min at room temperature. A 500 μL aliquot of the supernatant was incubated at 85 °C with 2 U of thermostable α-amylase for 90 min to fully hydrolyze starch into glucose (Moreno *et al*., 2010). The resulting soluble glucose concentration was determined using 3,5-dinitrosalicylic acid (Miller, 1959) in a microplate reader at 540 nm, using a glucose (Sigma - Aldrich, Sr. Louis, MO, USA) calibration curve (Shao and Lin, 2018). The soluble sugars concentrations were obtained according to González-Lemes *et al*. (González-Lemes *et al*., 2023). Samples were analyzed by HPLC (Waters-Millipore 510 HPLC) using a Hypersil Gold Amino carbohydrate analysis column (Thermo Scientific) at constant temperature (30 °C) using acetonitrile:water (3:1) as the mobile phase, with a flux of 1 mL min^−1^. Concentrations were obtained using calibration curves for sucrose, glucose, and fructose (Sigma-Aldrich).

### Plant water status parameters

Plant water status was determined by obtaining the relative water content (RWC) and osmotic potential (Ψs) from mature fully expanded leaves from six plants, per species and per treatment, at the mid-pod filling stage. RWC was obtained from three discs (1 cm diameter) per leaf, using six leaves per plant. Immediately after cutting, the discs were weighed to determine their fresh weight (FW). They were then incubated in distilled water at room temperature for 24 hours to get their turgid weight (TW). Following this, the discs were dried in an oven at 60 °C until a constant weight was reached (dry weight: DW). RWC was calculated using the following equation RWC = [(FW-DW)/(TW-DW)] x 100 (Rosales *et al*., 2013; Weatherley, 1950). The osmotic potential was obtained from three discs (1 cm diameter) per leaf, using the same number of leaves and plants as described above. After cutting, the discs were immediately frozen in liquid nitrogen and subjected to five cycles of freezing (1 min in liquid N_2_), thawed (10 min at 25°C) and then centrifuged for 20 min at 14,000 rpm. The osmolality of the sample was determined from 10 μL of the supernatant, with a vapor pressure osmometer (Wescor VAPRO Model 5600, Wescor, Inc., Logan UT USA). The values obtained in mmol/Kg were converted to MPa using the Van’t Hoff equation (Money, 1989).

### Leaf morphological analysis

The leaf area (LA) and specific leaf area (SLA) were obtained from 20 mature leaves that were initially tagged in an early developmental stage at the onset of flowering, when watering was halted, and were collected at the mid-stage of grain filling when leaves were fully expanded. The leaf samples were scanned to obtain their area using ImageJ. Thereafter, leaves were dried at 60 °C until weight was constant. SLA was calculated using the following equation SLA = LA (Leaf area) / DW (Dry weight) (Bray and Reid, 2002). Additionally, the total leaf biomass was obtained from ten plants from three independent experiments (from treatment and species), from a different set at the mid-stage of grain filling. Leaves were dried at 60 °C until weight was constant. From this set of experiments, the total number of leaves was also obtained.

### Light and transmission electron microscopy

For this analysis, we used fully expanded mature leaves from four plants, per species and per treatment, at the mid-stage of grain filling to distinguish possible differences in their anatomy. The cross sections, taken from the middle part of the leaves, were immediately fixed in 2.5% (v/v) glutaraldehyde (GTA) (Agar cat. R1020) and 4% (v/v) paraformaldehyde (PFA) (Electron Microscopy Sciences) in 0.1 M phosphate buffer (PB) pH 6.5 during 1 hour in a vacuum chamber, then they were fixed for 24 hours at 4°C. Afterwords, samples were rinsed with PB under dark and post-fixed with 1% (w/v) osmium tetroxide (Electron Microscopy Sciences) in PB for 2 hours at 4 °C. After this, samples were rinsed twice with PB and once with distilled water before gradual dehydration in a series of increasing ethanol concentrations (30, 50, 70, 80, 90, 95, and 100%) (20 min in each step). The dehydration mixture was gradually replaced with a mix of propylene oxide and epoxy resin as follows: 8 h in 3:1, 16 h in 1:1, and 6 h in 1:3, and subsequently embedded in epoxy resin (Epoxy Embedding Medium Kit resin, 45359-1EA-F Sigma-Aldrich) using silicone molds to polymerize in an oven (BOEKEL) at 60 °C for 48 h. Thereafter, semi-thin leaf sections (0.5 µm) were obtained in a Leica Ultracut R ultramicrotome (Leica Microsystems), which were stained in a 1% (w/v) toluidine blue (Sigma-Aldrich), 1% (w/v) sodium borate (Fermont) solution. The complete leaf cross sections were observed with a Nikon Eclipse E600, using 40X objective. Original light micrographs were recorded digitally with a Nikon Digital Sight 10 camera.

### Mesophyll cell type analysis

To obtain the proportion of the different mesophyll cell types from the leaf cross-sections, we labeled the cells in the digital images using Adobe Photoshop, then we binarized the images to determine the percentage of each cell type (palisade and spongy parenchyma and epidermal cells) (Supplementary Fig. S1A). For this analysis, we used 12 digital images, per species and per treatment, from four leaves. Based on the images taken from transverse sections of mature leaves, we analyzed different sections (155 μm in length) to quantify the density and width of palisade cells. Adobe Photoshop software was used for image alignment, while ImageJ was employed to extract the area for analysis and perform measurements. A total of seven images per treatment and species were analyzed to quantify cell density, and 20 cells per treatment and species were measured to determine their width. This parameter was defined as the value (in μm) at the point of greatest thickness of each cell. The proportion of spongy parenchyma cells was obtained by measuring the area occupied by these cells relative to the total area of the spongy parenchyma layer.

### Xylem vessel analysis

For this analysis, we focused on the vessels of the central vein, using semi-thin leaf transverse sections (0.5 µm) stained as described above. To obtain the total xylem area, we measured the area of the internal space within the visible discs of all the xylem vessels in each digital image of central veins. The sum of the areas of all central vein vessels was considered as the xylem total area per treatment and per species (Supplementary Fig. S1B). The area per xylem vessel was calculated by the mean of the xylem area per vessel, using 20 vessels.

For leaf venation analysis, we tagged 14 leaves, at early development, at the onset of flowering, when watering was halted. At the end of the treatment (mid-stage of grain filling), the completely expanded tagged leaves were harvested and circles of 1 cm in diameter were obtained from the middle part of the leaves, containing the central vein (mid-rib). The samples were immersed in 95% ethanol and glacial acetic acid solution in a 3:1 ratio for 48 – 72 h until complete clearing (Himes *et al*., 2021). Thereafter, samples were washed with distilled water and stained in 0.01% (w/v) safranine aqueous solution during 48 h (Garcia-Gutierrez *et al*., 2020). Images were examined in a stereoscopic microscope Nikon SMZ1270 (Nikon Instruments Inc.) and analyzed using ImageJ to measure the length of major veins, including first-, second-, and third-order veins (Supplementary Fig. S1C).

### Cuticle thickness analysis

For this study, ultrathin leaf cross sections placed on copper grids were stained for transmission electron microscopy (TEM) with 2.5% (v/v) uranyl acetate (EMS) for 20 min and 0.3% (v/v) lead citrate (EMS) for 10 min. The sections were rinsed with distilled water and dried to be observed at 80kV in a ZEISS Libra 120 plus electron microscope (Zeiss AG Oberkochen, Germany) with a GATAN ultrascan 1000 CCD camara (Gatan, Pleasanton, California, USA) coupled to a DigitalMicrograph. For this analysis, we examined 16 images (magnification 8000X) from three leaves of three plants, per species and treatment using the DigitalMicrograph^®^ software.

### RNA extraction

Total RNA was obtained from fully expanded trifolia from three plants, per treatment and per species, at the mid-stage of grain filling. After harvesting, leaves were immediately frozen in liquid nitrogen and kept at −70°C until use. RNA was extracted following the cetyltrimethylammonium bromide method as described by Acosta-Maspons *et al*. (Acosta-Maspons *et al*., 2019).

*Quality of the RNA-seq, read mapping to the P. vulgaris reference genome, and differentially transcript abundance*.

RNA integrity was verified using Agilent 2100 Bioanalyzer (Agilent Technologies, Inc.). For library construction and RNA sequencing, RNA from three independent replicates were sent to Laboratorio de Servicios Genómicos of Unidad de Genómica Avanzada (UGA), Irapuato, México, or to Unidad Universitaria de Secuenciación Masiva y Bioinformática of the Universidad Nacional Autónoma de México (UUSMB-UNAM), Cuernavaca, México. Samples were sequenced using DNBSEQTM system in a MGISEQ-2000 platform obtaining 22 millions of high-quality reads per library. Libraries were constructed from RNA samples obtained from three plants per treatment. The quality of the sequenced libraries was analyzed by FastQC v0.11.2 (www.bioinformatics.babraham.ac.uk/projects/fastqc/) and MultiQC v1.10.1 (Ewels *et al*., 2016). Adapters were removed using Cutadapt v3.2 (Martin, 2011) in a paired configuration, while to obtain high-quality reads (Quality Control >20) from libraries, we applied Trimmomatic v0.36 PE (Bolger *et al*., 2014) using the “KeepBothReads” option with a minimum length of 50 bp. Filtered reads of *Phaseolus vulgaris,* cultivar Pinto Saltillo, or *Phaseolus acutifolius* (line TARS-Tep 32) transcriptomes were mapped to the transcripts of the *P. vulgaris* v2.1 genome (*P. vulgaris* v2.1, DOE-JGI, and USDA-NIFA; http://phytozome.jgi.doe.gov/) with Bowtie2 v2.4.2 (Langmead and Salzberg, 2012) in the “very-sensitive” option. Tables of read counts were obtained from the same files with bowtie2 with Python scripts (https://github.com/genomica-fciencias-unam/tabla_genes/tree/english) (Romero *et al*., 2021). Correlations of the libraries from cultivars and conditions were correctly separated as *P. vulgaris* and *P. acutifolius*. Likewise, libraries from growth conditions, well-watered, and drought, were grouped between replicates in both *P. vulgaris* and *P. acutifolius*, respectively (Supplementary Fig. S4). Differential abundant transcripts (DAT) were addressed by DESeq2 (Love *et al*., 2014) in R v4.0 (https://www.r-project.org/), with p-value < 0.05 and Log2FoldChange > 1 or < −1 to identify significant differentially transcript abundance. Data exploration was performed with a principal component analysis (PCA) in R v4.0 (https://www.r-project.org/).

### Gene ontology enrichment analysis

Functional annotation and transcript enrichment analyses were performed using KEGG Orthology (KO) and Gene Ontology (GO) terms from the KEGG (https://www.kegg.jp/), Phytozome (http://phytozome.jgi.doe.gov/) and PANTHER (https://pantherdb.org/) databases (Thomas *et al*., 2003). Differentially abundant transcripts (DAT), identified either between transcripts accumulated in *P. vulgaris* and *P. acutifolius* under well-irrigated conditions, or between drought and control conditions within each species, were used for KEGG or GO term enrichment analysis, and GO terms with a *p*-value <0.05 were considered significantly enriched.

### Statistical analysis

Statistical analysis was performed by Student’s *t*-test or Mann-Withney U test using IBM SPSS Statistics v 21. Further details are included in the figure legends.

## Results

### Leaf water status during terminal drought is similar in the cultivars of both Phaseolus species

To evaluate the water status of the studied organs at the end of the terminal drought treatment during the mid-pod filling stage, we measured relative water content (RWC) (Fig. 1A) and osmotic potential (Ψs) (Fig. 1B) under well-watered, and water limited conditions. Additionally, we assessed stomatal conductance to complement these measurements (Fig. 1 C). At the time of measurement, all leaf samples exhibited a similar reduction in stomatal conductance, with a decrease of 76% in *Pv* and 72.5% in *Pa* (Fig. 1 C). This reduction was consistent with the maintenance of RWC in all leave samples, although *Pa* leaves showed a slightly but significant higher decrease in RWC in response to the stress treatment (Fig. 1A). The significant reduction in osmotic potential in the leaves of stressed plants from the cultivars of both species, with a 21.6% decrease in Pa and 10.85% in Pv, is consistent with the RWC values (Fig. 1B).

**Figure 1.**
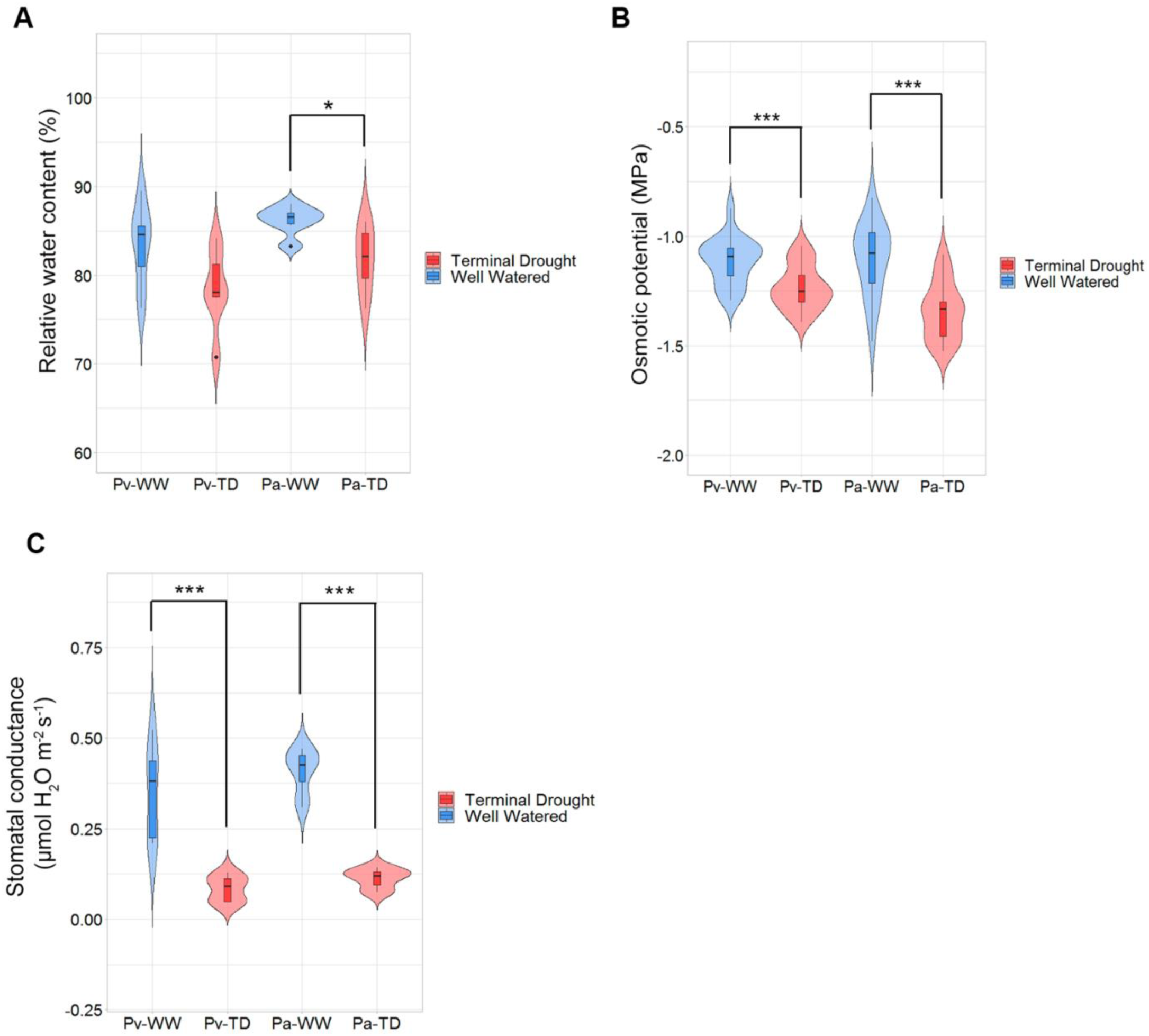
(A) Relative water content (RWC) expressed in (%) measured in leaves of *Phaseolus vulgaris* (*Pv*) and *Phaseolus acutifolius* (*Pa*) in well-watered (WW) and terminal drought (TD) conditions, bars represent the standard deviation, n=6. (B) Osmotic potential expressed in MPa measured in leaves of *Pv* and *Pa* in WW and TD conditions, bars represent the standard deviation, n=18. (C) Stomatal conductance expressed in µmol H_2_O m^−2^ s^−1^ measured in leaves of *Pv* and *Pa* in WW and TD conditions, bars represent the standard error, n=7. Significant differences were found using the *t*-test or Mann Whitney U test comparing between treatments. * *p* < 0.05 *** *p* < 0.001.

### Leaf venation structure in the leaves of P. acutifolius cultivar under terminal drought suggests an efficient water distribution

Considering the relevance of water distribution across leaf tissues for their functional maintenance, we evaluated possible anatomical modifications in the leaf venation system as a response to terminal drought of the studied *Phaseolus* cultivars. For this, we analyzed the major vein length density (MVLD) (mm/cm^2^), measuring MVL (mm) per area (cm^2^) of mature leaves (Fig. 2A, 2B, and Supplementary Fig. S1C). We found that *Pa* leaves exhibited a higher MVLD in stressed plants compared to well-watered samples (*p* < 0.001) (Fig. 2A). In contrast, no significative differences were observed for MVLD in *Pv* leaves between stress and non-stress conditions (Fig. 2A). Because vein density is related to leaf area, we determined this trait as well. The results showed that *Pa* leaves are significantly smaller than *Pv* leaves, with a mean leaf area of 20 cm^2^ in well-watered plants, compared to 48 cm^2^ in *Pv* leaves (Fig. 2C). As expected, plants subjected to water deficit treatment showed a significant reduction in leaf area, with *Pv* leaves showing a greater decrease (40%) than for *Pa* leaves (19.4%) (Fig. 2C). This finding aligns with the higher MVLD observed the smaller leaves of *Pa* under water limitation compared with well-watered conditions (Fig. 2D), in contrast to the decreasing MVLD trend seen in *Pv* under stress (Fig. 2D). These results suggest that leaf size and major vein density are coordinated traits that may underlie a more efficient water distribution and conservation strategy in the photosynthetic tissues of *Pa*.

**Figure 2.**
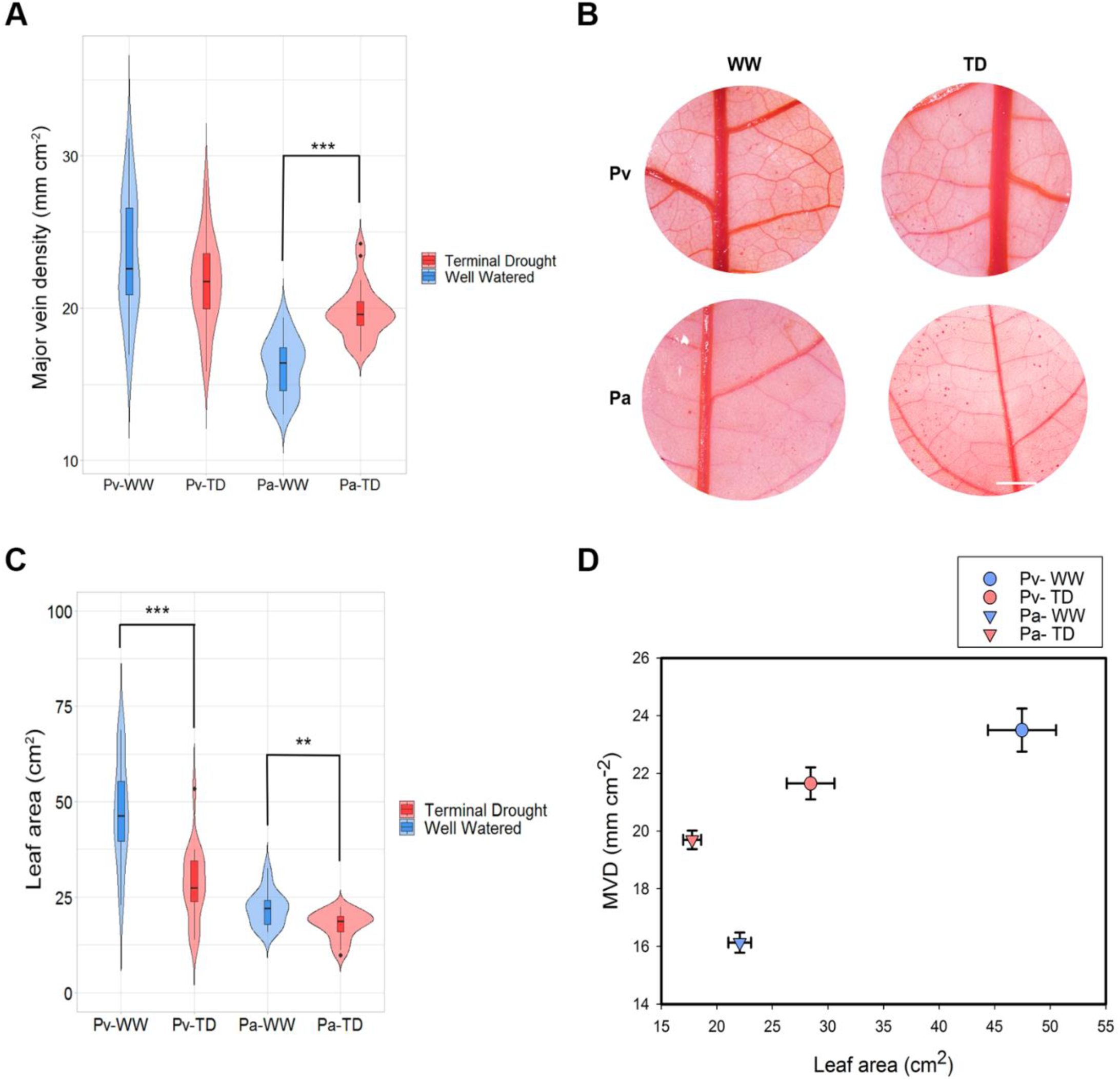
(A) Major vein length density (MVLD) expressed in (mm cm^−2^) measured in leaves of *Phaseolus vulgaris* (*Pv*) and *Phaseolus acutifolius* (*Pa*) in well-watered (WW) and terminal drought (TD) conditions, bars represent the standard deviation, n=28. (B) Images of the leaves cleared and stained with safranin of *Pv* and *Pa* in WW and TD conditions, the scale bar is equivalent to 2 cm. (C) Leaf area expressed in cm^2^ measured in leaves of *Pv* and *Pa* in WW and TD conditions, bars represent the standard error, n=20. (D) Comparison of the MVLD and leaf area. Significant differences were found using the *t*-test or Mann Whitney U test comparing between treatments. ** *p* < 0.01 *** *p* < 0.001.

Because the relevance of the xylem system in shielding plant water transport and distribution under water restriction (Ahmad *et al*., 2018; Huang *et al*., 2022; Nogueira *et al*., 2020), we also evaluated the total lumen area of xylem vessels in the central midrib (Fig. 3A and Supplementary Fig. S1B). This analysis revealed that this parameter was larger in *Pv* than in *Pa* leaves under non-stress conditions, although we did not find significant differences (Fig. 3B). Under stress conditions, the lumen area of the xylem vessels was similar in both species. *Pv* xylem total lumen exhibited a decreasing tendency (31.3 %) in response to stress, while for *Pa* no significant difference was found (Fig. 3B). When analyzing the lumen area of individual xylem vessels, we noticed that the xylem vessels are narrower in response to drought (Fig. 3C). Moreover, the *Pv* xylem showed a smaller number of vessels with smaller lumen area under terminal drought treatment than *Pa* xylem vessels (Fig. 3C and 3D), supporting that *Pa* xylem system favors a more efficient water distribution.

**Figure 3.**
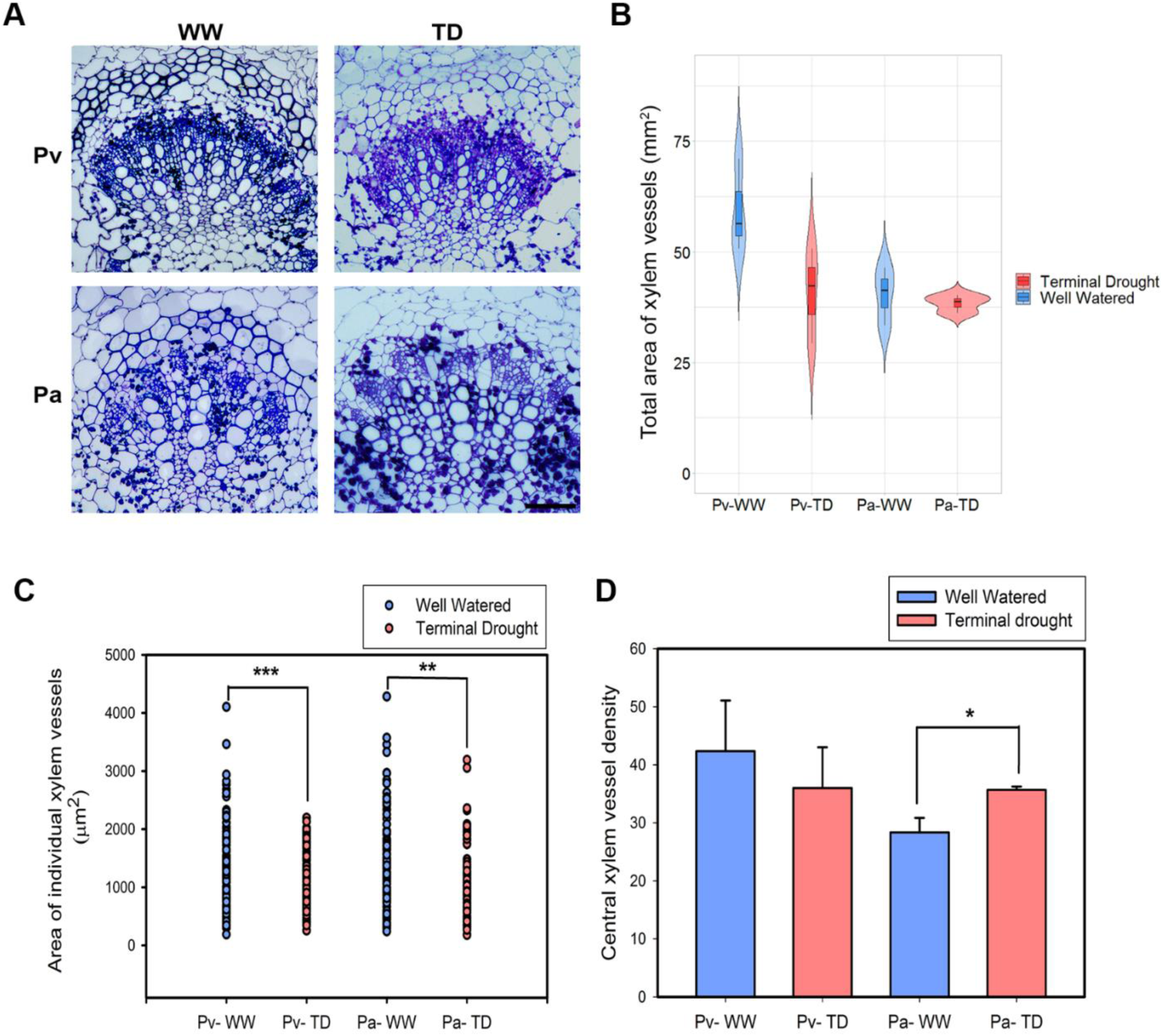
(A) Images of cross sections of the midrib stained with toluidine blue where the xylem vessels can be observed. the bar is equivalent to 40 µm. (B) Total area of xylem vessels expressed in (mm^2^) measured in leaves of *Phaseolus vulgaris* (*Pv*) and *Phaseolus acutifolius* (*Pa*) in well-watered (WW) and terminal drought (TD) conditions, bars represent the standard deviation, n=3. (C) Area of individual xylem vessels expressed in µm^2^ measured in leaves of *Pv* and *Pa* in WW and TD conditions, n=127, 108, 85, 107. (D) Central xylem vessel density, bars represent the standard deviation, n=3. Significant differences were found using the *t*-test or Mann Whitney U test comparing between treatments. * *p* < 0.05 ** *p* < 0.01 *** *p* < 0.001.

### Leaf cuticle thickness increases in the leaves of P. acutifolius cultivar in response to terminal drought

The cuticle is an extracellular hydrophobic protective layer covering the epidermis of leaves and other aerial parts of the plant. Along with stomatal function, the cuticle plays a key role in minimizing leaf water loss, making it essential for protecting leaves from dehydration during water deficit conditions (Chen *et al*., 2020; Yeats and Rose, 2013). To assess the involvement of this structure in *Phaseolus* leaves under terminal drought, we analyzed images obtained through TEM to evaluate potential changes in leaf cuticle thickness (Fig. 4A). The results revealed a significant reduction (*p* < 0.001) in the thickness of the adaxial cuticle in *Pv* leaves under stress, whereas *Pa* leaves showed a slight but significant increase (*p* < 0.05) in this trait (Fig. 4B).

**Figure 4.**
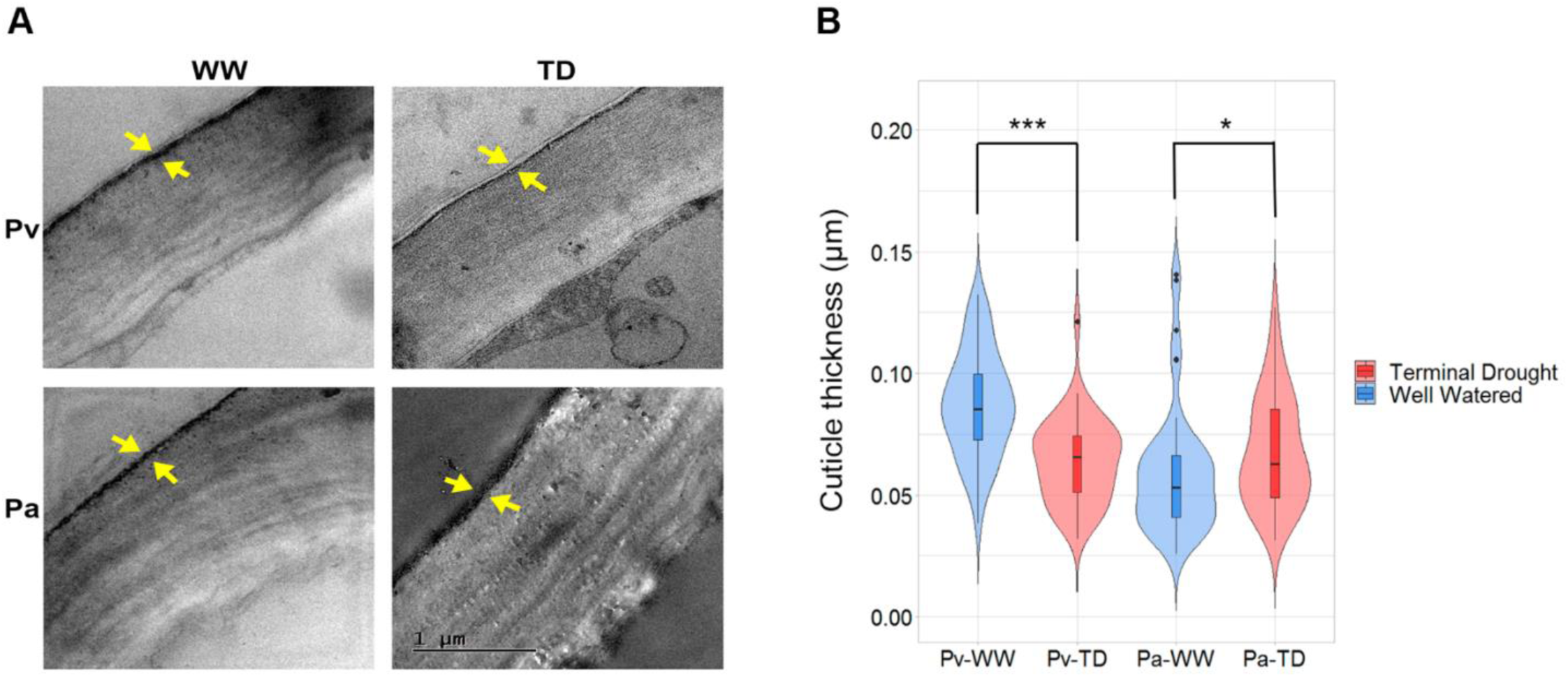
(A) Images of the adaxial epidermis taken with a transmission electron microscope where the cuticle of the leaves can be seen. (B) Thickness of the cuticle expressed in (µm) measured in leaves of *Phaseolus vulgaris* (Pv) and *Phaseolus acutifolius* (Pa) in well-watered (WW) and terminal drought (TD) conditions, bars represent the standard error, n=48. Significant differences were found using the *t*-test or Mann Whitney U test comparing between treatments * *p* < 0.05 *** *p* < 0.001.

### Terminal drought treatment affects leaf photosynthetic activity in cultivars of both Phaseolus species

To assess the relationship between leaf anatomical traits and photosynthetic performance in *Pv* and *Pa* plants under water-limiting conditions during the reproductive stage, we first evaluated the net CO_2_ assimilation rate (A_n_) and intercellular CO_2_ concentration (C_i_) in mature leaves. Interestingly, under well-watered conditions, *Pa* exhibited higher A_n_ values (20.79 μmol CO_2_ m^−2^ s^−1^) at lower intercellular CO_2_ concentrations (252.48 μmol CO_2_ mol air^−1^) compared to *Pv* net photosynthesis (13.68 μmol CO_2_ m^−2^ s^−1^) at a higher intercellular CO_2_ level (287.17 μmol CO_2_ mol air^−1^) (Fig. 5A). In contrast, under terminal drought, both species showed a reduction in A_n_. Although *Pa* showed slightly higher A_n_ values than *Pv*, the decline was more pronounced in *Pa* (9.73 μmol CO_2_ m^−2^ s^−1^) compared to *Pv* (7.80 μmol CO_2_ m^−2^ s^−1^) (Fig. 5A).

**Figure 5.**
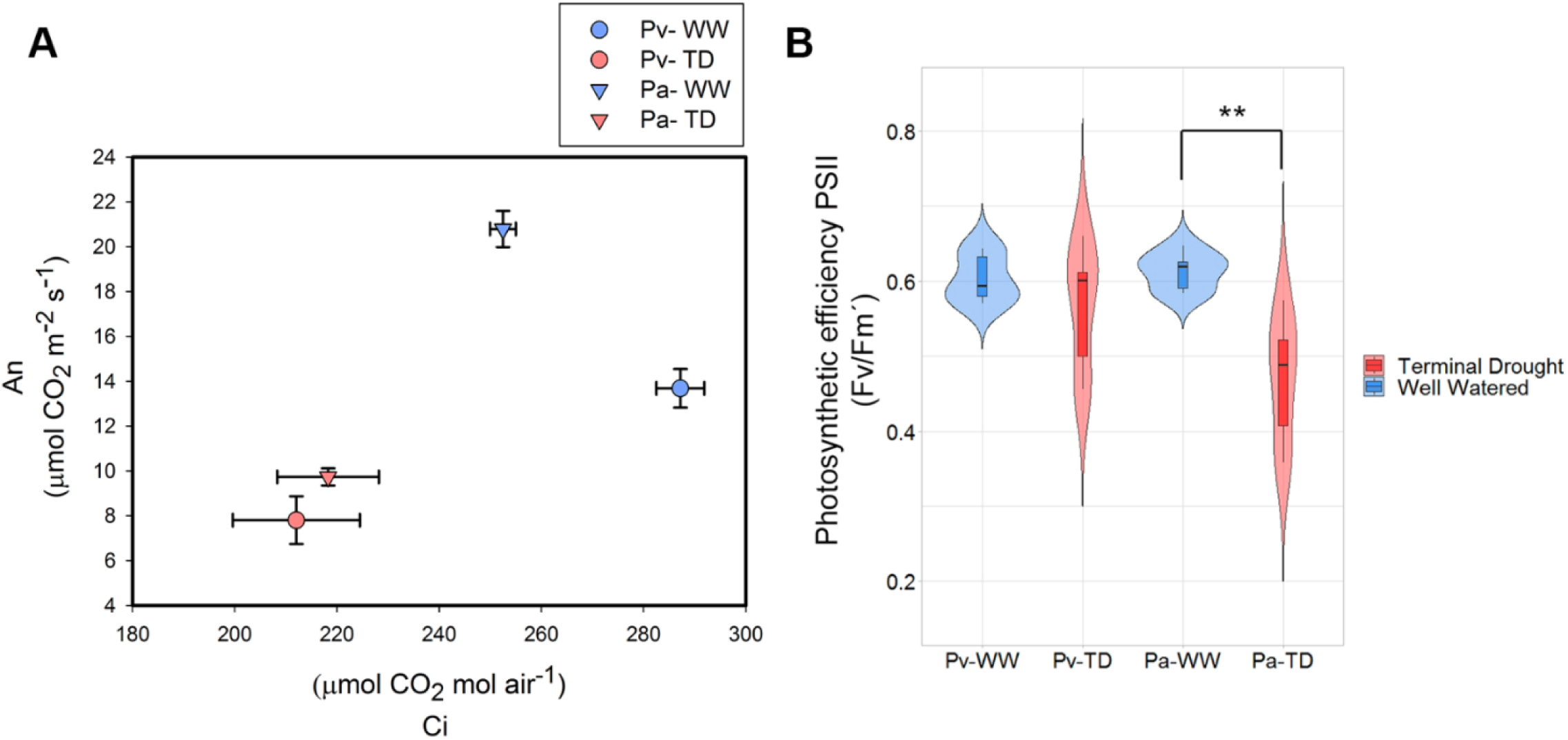
(A) Net photosynthesis rates (A_n_) and intercellular CO_2_ concentration (C_i_) measured in leaves of *Phaseolus vulgaris* (*Pv*) and *Phaseolus acutifolius* (*Pa*) in well-watered (WW) and terminal drought (TD) conditions, bars represent the standard error, n=7. (B) Photosynthetic efficiency PSII expressed in (Fv/Fm’) measured in leaves of *Pv* and *Pa* in WW and TD conditions, bars represent the standard error, n=6. Significant differences were found using the *t*-test or Mann Whitney U test comparing between treatments. ** *p* < 0.01.

To evaluate whether these observations were related to changes in photosystem II efficiency, we measured this parameter and found that, in agreement with the reduction in net photosynthesis and C_i_, *Pa* leaves also experienced a greater decline in this parameter under stress (Fig. 5B). Nevertheless, we did not find significant differences between the two species under well-watered conditions (Fig. 5B). These data agree with the chlorophyll content obtained for both genotypes under the two experimental conditions tested (Supplementary Fig. S2).

### P. acutifolius cultivar exhibits a larger efficiency of carbon fixation

The high net CO_2_ assimilation rate values under relatively low CO_2_ concentration inside the leaf air space (C_i_) exhibited by *Pa* (Fig. 5A) prompted us to evaluate the response of photosynthesis to different C_i_ values. For this, we conducted A_n_ measurements as a function of C_i_ (A_n_/C_i_ curves) (Supplementary Fig. S3). These curves allow the calculation of parameters such as the maximum carboxylation rate (V_cmax_), which measures the efficiency of carbon fixation by Rubisco (ribulose.1,5-biphospahte carboxylase/oxygenase) (Bernacchi *et al*., 2013).

When comparing A_n_ values between treatments, the cultivars of both species showed a significant decrease (*p*<0.05) under water deficit (Fig. 6A). Of note, as CO_2_ concentration increased, *Pa* under drought exhibited a response trend similar to that of *Pv* under well-watered conditions, indicating a more efficient CO_2_ assimilation capacity in *Pa*. This effect became more evident when comparing between species, where *Pa* consistently displayed significantly higher A_n_ values by increasing C_i_ levels (Fig. 6A), showing an even greater CO_2_ fixation efficiency under optimal irrigation. A similar trend was observed in V_cmax_. Although no significant differences were found between treatments (Fig. 6B), *Pa* showed a V_cmax_ 1.2 times higher than *Pv* under optimal irrigation (*p*<0.01) (Fig. 6B), confirming its greater carboxylation efficiency.

**Figure 6.**
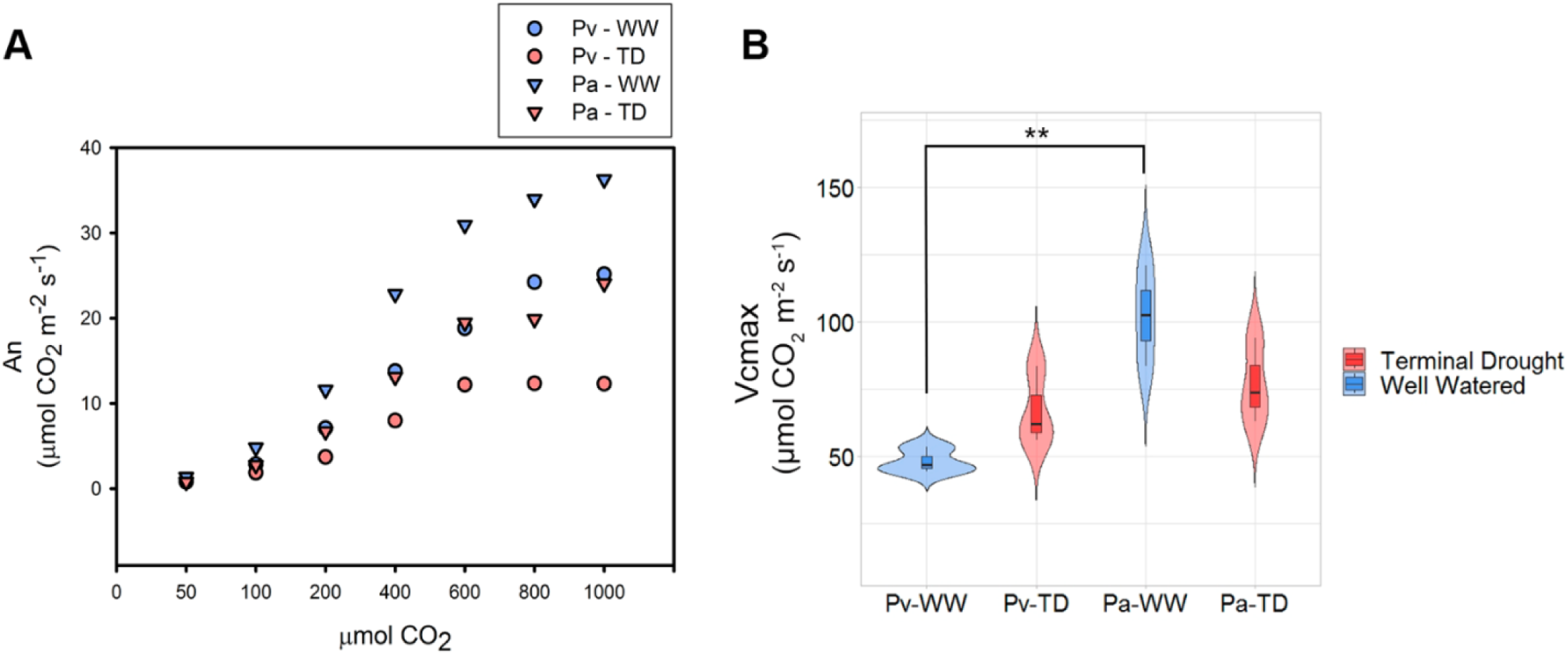
(A) CO_2_ response (An/Ci) curves measured in leaves of *Phaseolus vulgaris* (*Pv*) and *Phaseolus acutifolius (Pa*) in well-watered (WW) and terminal drought (TD) conditions, n=3. (B) Photosynthetic efficiency PSII expressed in (µmol CO_2_ m^−2^ s^−1^) measured in leaves of *Pv* and *Pa* in WW and TD conditions, bars represent the standard deviation, n=3. Significant differences were found using the *t*-test or Mann Whitney U test comparing between treatments. ** *p* < 0.01.

### Leaf air space and palisade parenchyma cell density and thickness in both Phaseolus species under terminal drought

Since V_cmax_ depends on the amount of CO_2_ reaching the chloroplasts (Manter and Kerrigan, 2004; Niinemets *et al*., 2009), the observed differences may reflect variations in palisade cell anatomy and/or leaf intercellular air space between *Pa* and *Pv* leaves (Evans, 2021). This size of palisade parenchyma cells is known to affect the chloroplasts surface area exposed to intercellular spaces, CO_2_ diffusion efficiency, and ultimately, photosynthetic capacity (Egesa *et al*., 2024). To investigate this, we measured palisade cell traits using transverse leaf sections (Fig. 7A and Supplementary Fig. S1A). Our results showed no significant differences in palisade cell length (μm) between well-watered and drought-treated *Pa* leaves (Fig. 7B). In contrast, *Pv* leaves showed significantly longer palisade cells under drought conditions compared to well-irrigated plants (Fig. 7B). Regarding cell width, *Pv* showed a significant increase (*p* < 0.001) under drought, while *Pa* did not exhibit statistically significant changes between treatments (Fig. 7C). Nevertheless, when comparing the two *Phaseolus* cultivars, *Pa* consistently showed narrower palisade cells under both irrigation regimes (*p* <0.001) (Fig. 7C). As expected, this narrower cell structure was associated with a higher palisade cell density in *Pa* leaves compared to *Pv* (Fig. 7C). Regarding intercellular air space measurements, no differences were observed between the two species under well-watered conditions (Fig. 7D). In *Pv* leaves, intercellular air space remained unchanged between treatments, whereas in *Pa* leaves, a significant increase was detected in response to drought, favoring CO_2_ diffusion and uptake (Fig. 7D). We also quantified the total area of spongy parenchyma cells and found no significant differences between species or treatments (Supplementary Fig. S4).

**Figure 7.**
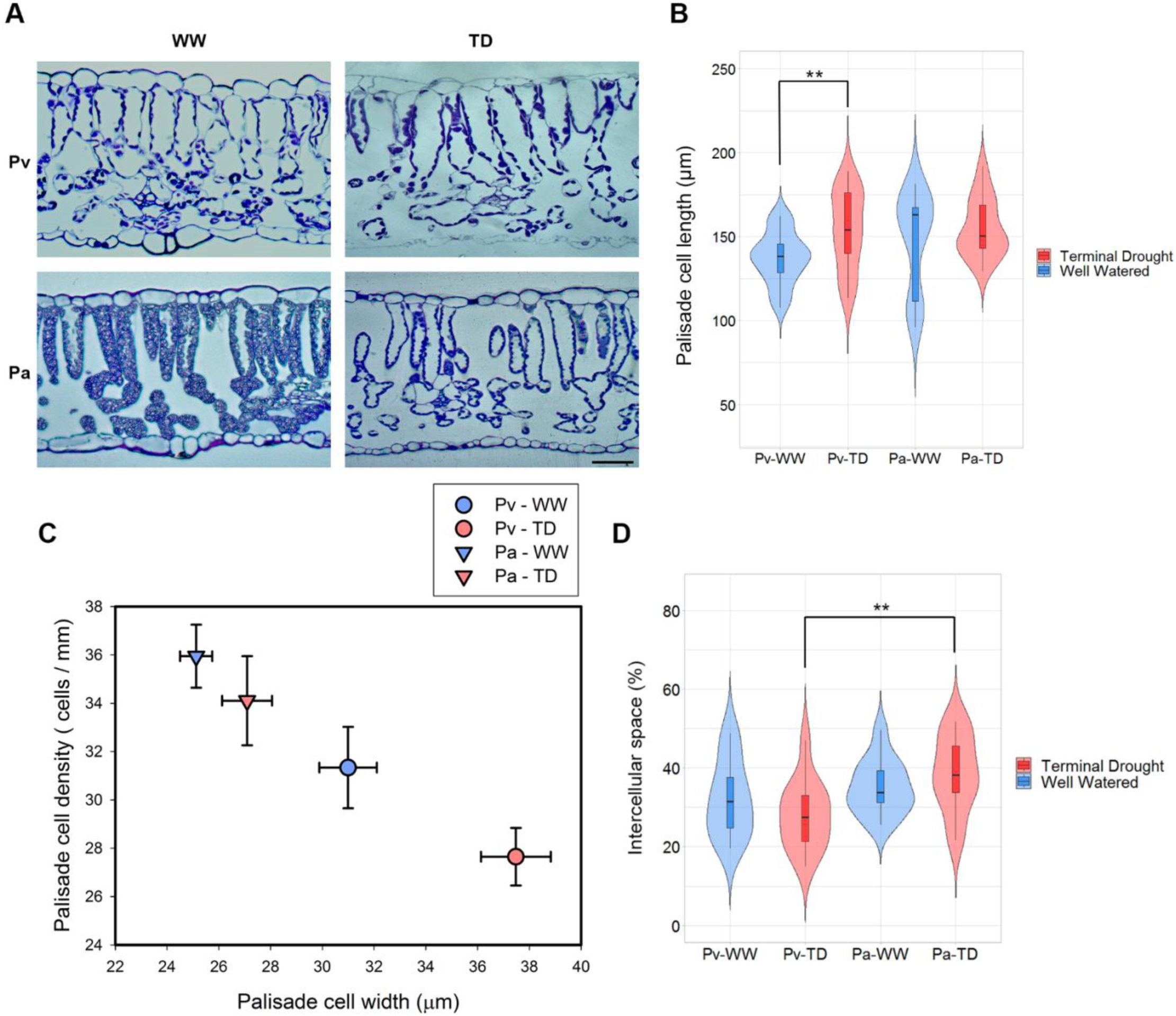
(A) Images of leaf cross-sections stained with toluidine blue (left) bar= 20 µm and (B) palisade cell length (right) expressed in (µm) measured in leaves of *Phaseolus vulgaris* (*Pv*) and *Phaseolus acutifolius* (*Pa*) in well-watered (WW) and terminal drought (TD) conditions, bars represent the standard deviation, n=24. (C) Palisade cell density expressed in (cells/mm) and palisade cell width expressed in (µm) measured in leaves of *Pv* and *Pa* in WW and TD conditions, bars represent the standard error, n=7, 20. (D) Percentage of intercellular space measured in leaves of *Pv* and *Pa* in WW and TD conditions, bars represent the standard error, n=12. Significant differences were found using the *t*-test or Mann Whitney U test comparing between treatments. ** *p* < 0.01.

### Changes in starch and sugar levels in response to terminal drought in both Phaseolus species

Starch plays a dual role as a product and a regulator of photosynthesis, functioning simultaneously as a carbon source and sink. In leaves, it is broken down to provide carbon for growth and development, while in various plant organs, it also serves as an important storage compound (AbdElgawad *et al*., 2020; Krasensky and Jonak, 2012; Long and Adams, 2023; MacNeill *et al*., 2017). Therefore, carbon fixation efficiency may be reflected in the amount of starch accumulated in leaves under well-watered and terminal drought conditions. To assess this, we measured the starch content in *Pv* and *P*a leaves under both treatments (Fig. 8A). Starch levels significantly decreased in both species under drought (*p* < 0.05 for *Pv* and *p* < 0.01 for *Pa*) (Fig. 8A). However, *Pa* showed consistently higher starch concentrations than *Pv,* 40% higher under irrigation and 88% higher under drought (Fig. 8A), suggesting that *Pa* maintains higher photosynthetic activity under both conditions.

**Figure 8.**
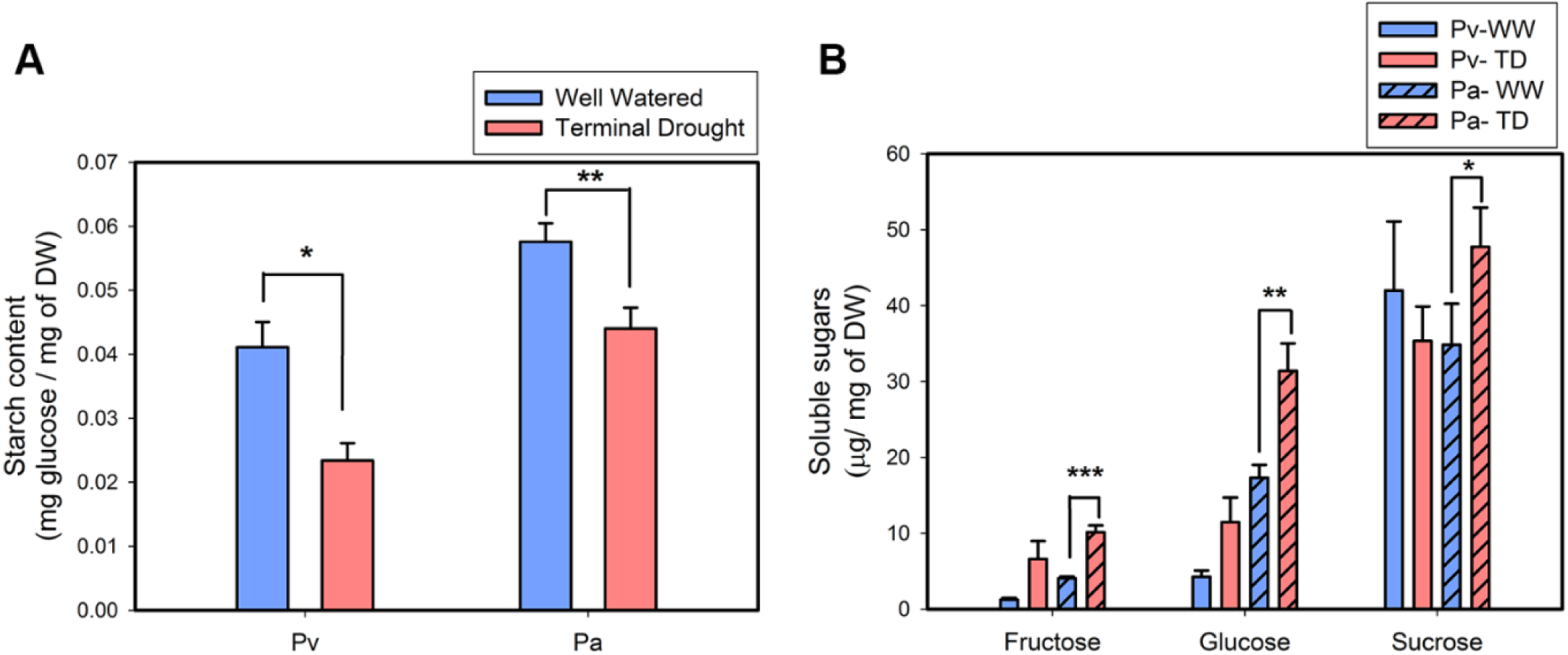
(A) Starch content expressed in (mg glucose/ mg of dry weight) measured in leaves of *Phaseolus vulgaris* (*Pv*) and *Phaseolus acutifolius* (*Pa*) in well-watered (WW) and terminal drought (TD) conditions, bars represent the standard error, n=27. (B) Fructose, glucose and sucrose expressed in µg/mg of dry weight measured in leaves of *Pv* and *Pa* in WW and TD conditions, bars represent the standard error, n=6-9. Significant differences were found using the *t*-test or Mann Whitney U test comparing between treatments. * *p* < 0.05 ** *p* < 0.01 *** *p* < 0.001.

Because sucrose is another key product of photosynthesis and the primary form of sugar used to transport energy for growth and storage (Everard, 2017), we also measured sucrose levels in both *Phaseolus* species under both treatments (Fig. 8B). Unlike starch content, sucrose levels significantly increased in *Pa* in response to drought, whereas no change was observed in *Pv*, further supporting the idea of a more efficient carbon fixation in *Pa*. In addition, we quantified glucose and fructose, the products of sucrose degradation, which are essential for energy supply, the synthesis of various metabolites and cellular structures, and the regulation of carbon flow (Braun *et al*., 2014; Ruan *et al*., 2010). Similar to sucrose, *Pa* showed a significant increase in glucose and fructose levels under drought, while no significant changes were detected in *Pv* under the same conditions (Fig. 8B). The comparable sucrose levels but lower concentrations of its degradation products in *Pv* compared to *Pa* suggest either more active sucrose degradation in *Pa* or a higher rate of sucrose synthesis in *Pv*, among other possible explanations.

### Terminal drought affects specific leaf area and aerial biomass

Differences in sugar storage and synthesis could influence resource allocation and shoot growth. To assess the impact of drought treatment on overall plant growth in the analyzed species, we measured the specific leaf area (SLA), a functional trait that represents the light-capturing surface area per unit of dry biomass and thus reflects carbon gain and plant growth (Legner *et al*., 2014; Wellstein *et al*., 2017). Measurements taken from fully expanded leaves showed that *Pv* plants under optimal irrigation had a higher SLA than those under water-limited conditions (Fig. 9A). In contrast, the lower SLA under water shortage corresponds to leaves with smaller surface area and higher density, resulting in a reduced transpiring surface. For *Pa* leaves under optimal irrigation, we observed a lower SLA compared to *Pv* leaves, consistent with their smaller, denser structure (Fig. 9A). Interestingly, under water deficit, the SLA of *Pa* leaves increase, suggesting that *Pa* favors a larger leaf area with lower density (thinner leaves) to cope with reduced water availability (Fig. 9A). This strategy may enhance the plant’s capacity to capture light under these adverse conditions. Additionally, to complement these findings, we measured the total dry leaf weight of both species under both conditions. Dry weight was assessed at the mid-grain filling stage, revealing a significant reduction in aboveground foliar biomass accumulation caused by drought treatment in both species (p < 0.001) (Fig. 9B). However, the biomass reduction was more pronounced in *Pv* (50%) than in *Pa* (35%), with Pa maintaining significantly higher dry leaf weight (p < 0.01) compared to *Pv* (Fig. 9B).

**Figure 9.**
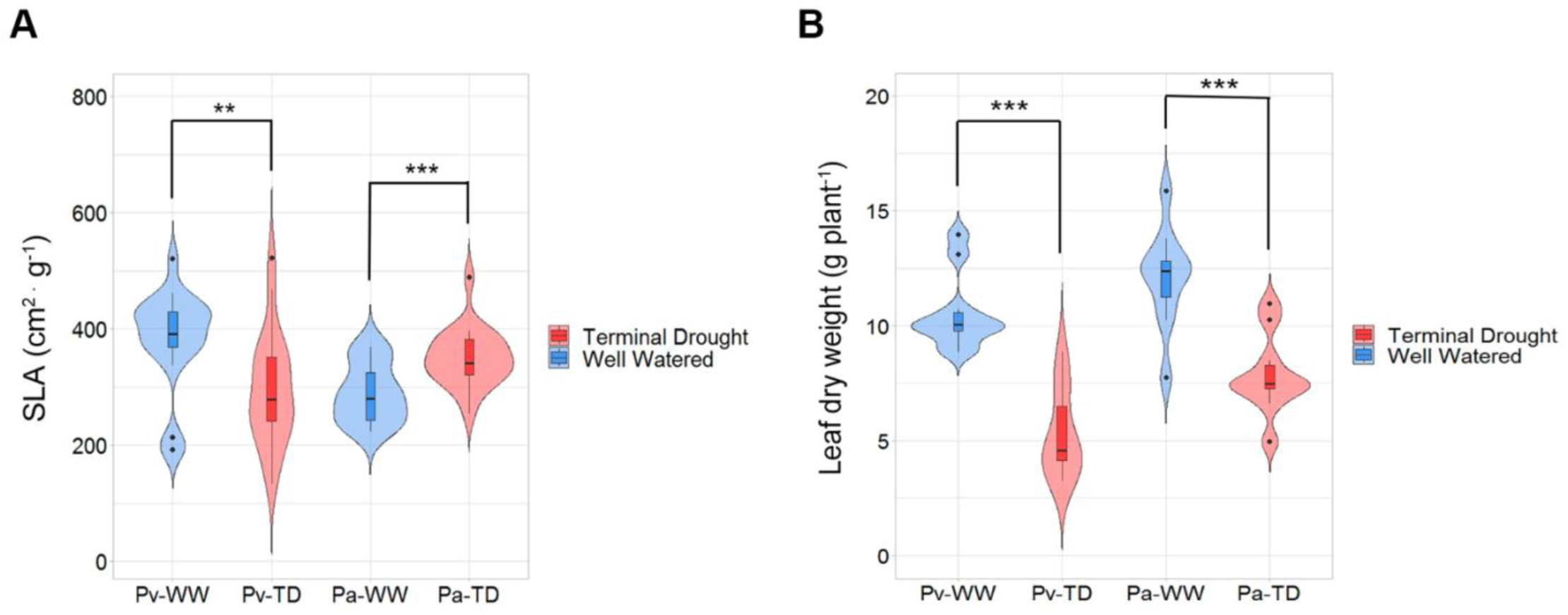
(A) Specific leaf area expressed in (cm^2^/g) measured in leaves of *Phaseolus vulgaris* (*Pv*) and *Phaseolus acutifolius* (*Pa*) in well-watered (WW) and terminal drought (TD) conditions, bars represent the standard error, n=21. (B) Leaf dry weight expressed in g per plant of *Pv* and *Pa* in WW and TD conditions, bars represent the deviation error, n=10. Significant differences were found using the *t*-test or Mann Whitney U test comparing between treatments. ** *p* < 0.01 *** *p* < 0.001.

### Transcriptomic analysis shows distinct responses between two Phaseolus cultivars

To gain deeper insight into the leaf traits of these species under terminal drought, we generated transcriptomes from leaves of *Pa* and *Pv* plants grown under well-watered and terminal drought conditions (see Materials and methods for details). As an initial step, we compared transcript abundance in each species under well-watered conditions and identified 3,654 and 3,453 DATs in *Pv* and *Pa* leaves, respectively (Fig. 10A).

**Figure 10.**
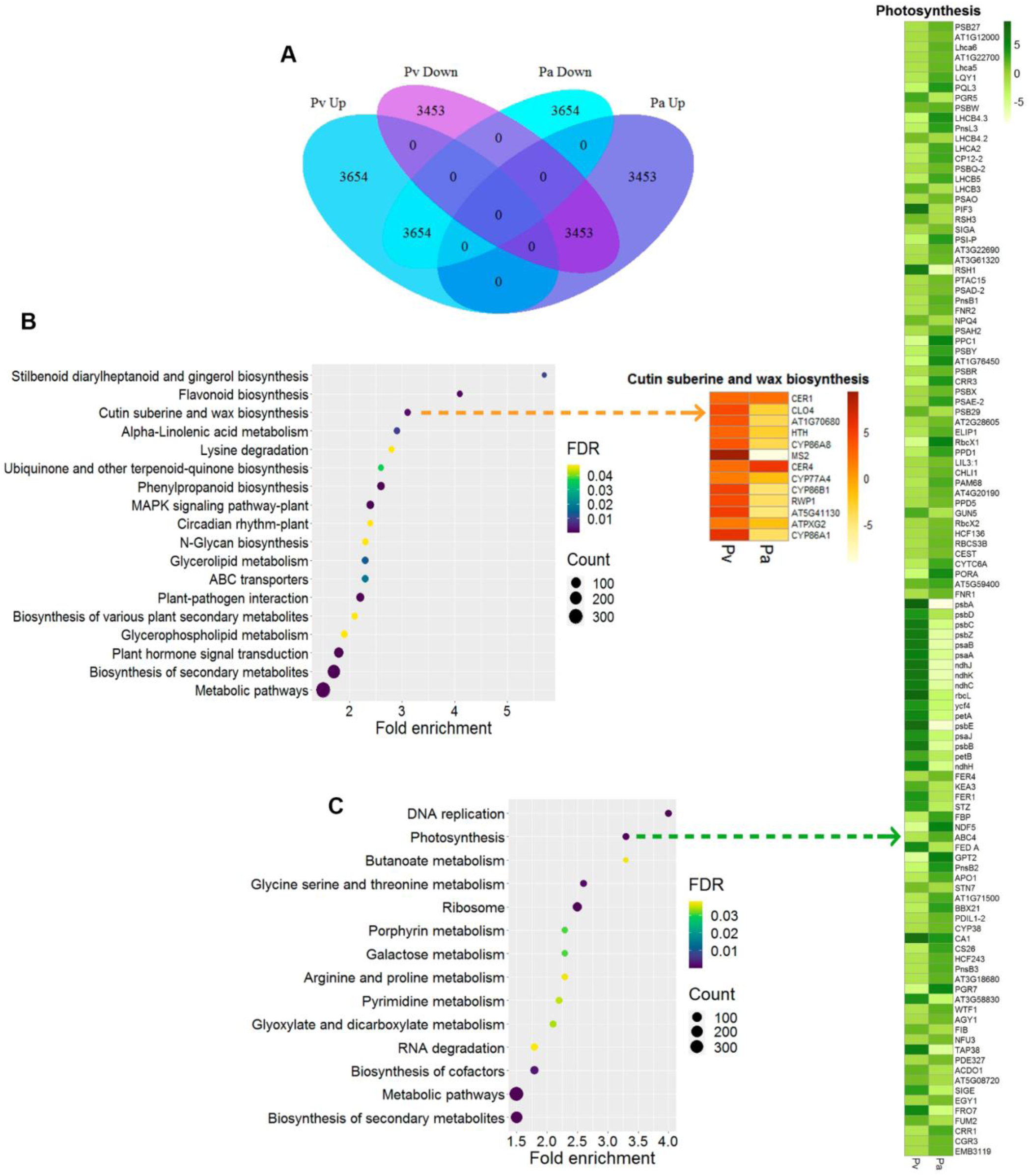
Transcriptomic analysis of *Phaseolus vulgaris* (*Pv*) and *Phaseolus acutifolius* (Pa) DATs from leaves under well-watered conditions. (A) Venn diagram showing the number of up- and down-regulated DATs from *P. vulgaris* and *P. acutifolius* leaves under well-watered conditions. (B) Enrichment analysis of *P. acutifolius* up-regulated DATs according to KEGG pathway classification, the heat map to the right shows the comparative abundance of *Pv* and *Pa* transcripts involved in the photosynthetic process. (C) Enrichment analysis of *P. vulgaris* up-regulated DATs according to KEGG pathway classification, the heat map to the right highlights the comparative abundance of *Pv* and *Pa* transcripts involved in cutin, suberin and wax biosynthesis. The color intensity represents the difference in transcript abundance, based on the log_2_fold change values.

KEGG pathway enrichment analysis showed that in both species, significantly enriched up-regulated transcripts were mainly associated with metabolic pathways, biosynthesis of secondary metabolites, and amino acid metabolism (Fig. 10B and C), consistent with their active growth under these conditions. Nevertheless, GO classification also identified stress-response transcripts, particularly in *Pv*, suggesting that this cultivar is either more sensitive to environmental fluctuations inherent to greenhouse conditions or activates a safeguard defense response (Supplementary Fig. S6A). In addition, *Pv* displayed enrichment of up-regulated transcripts linked to MAPK signaling, ABC transporters, plant hormone signal transduction, and cutin, suberin and wax biosynthesis (Fig. 10B). By contrast, *Pa* showed enrichment of transcripts involved in photosynthesis, biosynthesis of cofactors, DNA replication, and ribosome processes (Fig. 10C). These results are consistent with GO terms like lignin biosynthetic process and inorganic ion transmembrane transport for *Pv* (Supplementary Fig. S6B), and chloroplast organization, photosynthesis and light reactions, chloroplast RNA processing, and starch metabolism for *Pa* (Supplementary Fig. S6B). Considering the physiological and cellular traits of these cultivars, the enrichment of transcripts related to cutin and wax biosynthesis in *Pv* aligns with the thicker cuticle observed in *Pv* leaves compared with *Pa* under well-watered conditions (Fig. 4A and B). Likewise, the enrichment of photosynthesis and chloroplast processes related transcripts in *Pa* corresponds to the higher photosynthetic efficiency measured in *Pa* leaves from plants grown under non-stress conditions (Fig. 5B).

Analysis of the DAT under terminal drought identified 2,773 transcripts in *Pv* leaves, including 1,231 up-regulated and 1,542 down-regulated (Fig. 11A and Supplementary Fig. 5A and 5C). In *Pa* leaves 749 transcripts were detected, with 246 up-regulated and 503 down-regulated (Fig. 11A, Supplementary Fig. 5A and 5D). Both species shared 129 up-regulated and 157 down-regulated transcripts (Fig. 11A). The relatively. small number of DATs identified in *Pa* (749) and the limited overlap with *Pv* (286) (Fig. 11A) indicate that *Pa* mounts a more restrained transcriptional response to the drought treatment, consistent with its domestication in arid and semi-arid regions. When stress responsive transcripts were grouped, most were among the 129 commonly up-regulated transcripts, all of which exhibited higher accumulation levels in *Pv* than in *Pa* (Supplementary Fig. S8 and Supplementary Table S1). This pattern suggests that *Pa* may cope with drought stress by employing strategies that avoid water limitation, such as anatomical adjustments including greater major vein length, higher central xylem vessel density, and a thicker leaf cuticle, rather than through extensive transcriptional reprogramming. Because only 117 transcripts were up-regulated in *Pa*, KEGG pathway analysis grouped them exclusively into ‘secondary metabolite biosynthesis’ (Supplementary Table S2). This category actually encompasses transcripts involved in diverse processes, including cutin and wax biosynthesis, abscisic acid biosynthesis, sucrose and starch metabolism, carbohydrate metabolism, lipid metabolism, and photorespiration, among others (Supplementary Table S2). GO classification revealed transcripts linked to additional processes, from which underscore those related to abscisic acid and water deficit responses, as well as to cell wall, metabolism and biogenesis, development and morphogenesis processes (Fig. 11B and 11C).

**Figure 11.**
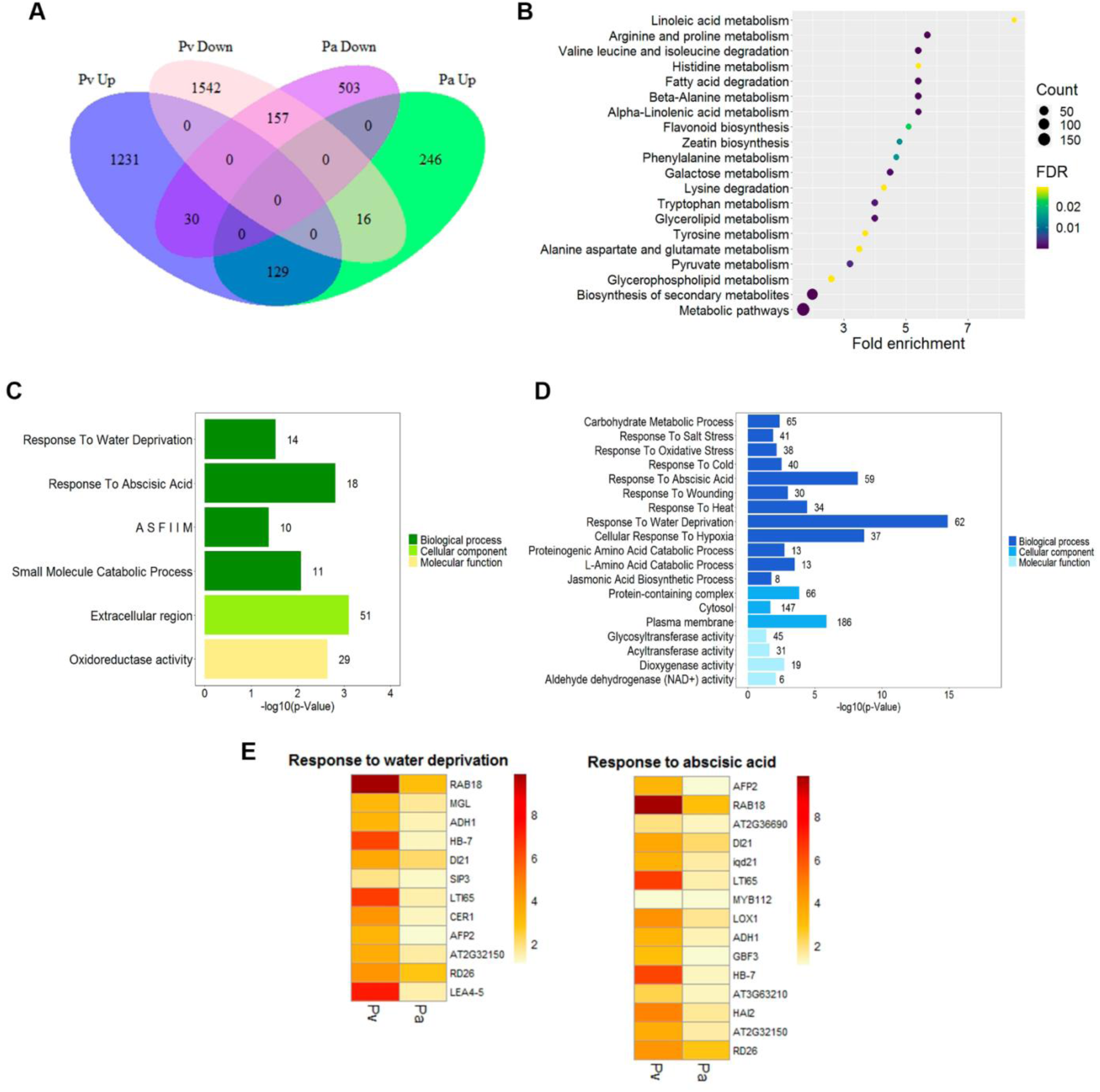
Comparative transcriptomic analysis of *Phaseolus vulgaris* and *Phaseolus acutifolius* DATs from leaves under terminal drought conditions. (A) Venn diagram showing the number of up- and down-regulated DATs from *P. vulgaris* and *P. acutifolius* leaves under terminal drought conditions. (B) Enrichment analysis of *P. vulgaris* up- regulated DATs according to KEGG pathway classification in response to terminal drought. (C) Representation of GO enrichment analysis of the up-regulated DATs of *P. acutifolius* in response to terminal drought. (D) Representation of GO enrichment analysis of the up-regulated DATs of *P. vulgaris* in response to terminal drought. (E) Heat maps showing the comparative abundance of water deficit and abscisic acid responsive transcripts. The color intensity represents the difference in transcript abundance based on log_2_fold change values.

Regarding transcriptomic response of *Pv* leaves, KEGG enrichment analysis revealed transcripts associated with diverse metabolic pathways, including secondary metabolite biosynthesis, alpha-linolenic acid metabolism, carbon metabolism (galactose and pyruvate), and the metabolism of several amino acids (Fig. 11D and E). GO analysis further highlighted the enrichment of stress-responsive transcripts, including those encoding different LEA proteins, molecular chaperones, small heat shock proteins, abscisic acid-responsive proteins, and antioxidant enzymes, among others (Fig. 11E). Many of these transcripts fall within the secondary metabolite biosynthesis category in the KEGG classification (Supplementary Table S2). In addition, transcripts involved in transport and plasma-membrane turnover were enriched, consistent with the KEGG identification of glycerolipid metabolism and possibly linked to the increased length of *Pv* palisade cells observed under drought.

## Discussion

Climate change is increasing the prospect and severity of droughts worldwide, including a rising frequency of terminal drought events (Yuan *et al*., 2023). This condition can have devastating effects on the cultivation of most crops, particularly those grown under rainfed conditions. Terminal drought severely impacts seed and fruit yield and quality, with far-reaching consequences for regional economies, food security, and public health. Because common bean, one of the most widely consumed legumes (Broughton *et al*., 2003; Gepts *et al*., 2008), is mostly grown under rainfed conditions, approximately 70% of its global production is affected by terminal drought, making it a major limiting factor (Beebe *et al*., 2013; Debouck *et al*., 2024). Due to its natural adaptation to arid climates (Bolger *et al*., 2014; Chen, 2024; Debouck *et al*., 2024), the tepary bean possesses unique traits that confer high drought and heat tolerance. Its broad adaptability and close genetic relation to common bean have drawn increasing attention as both a valuable genetic resource for improving common bean and a promising edible legume.

### Tepary bean exhibits drought-induced anatomical modifications to preserve leaf functionality

To get insights into the traits underlying the resistance to terminal drought in common and tepary beans, in this work, we characterized different leaf attributes of the common bean cultivar Pinto Saltillo, a genotype developed for high yield under terminal drought (Sanchez-Valdez *et al*., 2004), and of the tepary bean improved cultivar TARS-Tep32, known for its resistance to high temperatures and terminal drought (Porch *et al*., 2013; Porch *et al*., 2017). Under terminal drought treatment, although the leaves of both cultivars maintain their RWC, the osmotic potential in tepary leaves showed a considerably greater reduction (21.6%) compared to common bean (10. 85%). This indicates a more effective osmotic adjustment in tepary leaves, allowing it to better maintain cell turgor and support continued growth under water-limited conditions (Fang and Xiong, 2015; Ozturk *et al*., 2021). To identify anatomical attributes in the leaves of these genotypes that may contribute to minimizing water loss and enhancing water distribution efficiency under terminal drought, we also analyzed the morphology of major leaf veins. Our analysis revealed that tepary bean exhibits an increased density of major veins within a smaller leaf area compared to common bean. This trait may help compensate the delayed stomatal closure observed in tepary bean under terminal drought (Polania *et al*., 2022), potentially contributing to reduction in hydraulic vulnerability (Scoffoni *et al*., 2011). Similar strategies have been documented in other drought-adapted species, including *Caragana korshinskii, Helianthus annus, Gossypium hirsutum*, among others (Earley *et al*., 2024; Lei *et al*., 2018; Li *et al*., 2022). Although an increase in vein density is generally not be expected under stress conditions due to the requirement of enough resources (Beerling and Franks, 2010; Liao *et al*., 2023; Zhang *et al*., 2020), this observation suggests that tepary bean may retain sufficient reserves to invest in such structural modifications even under adverse environmental conditions. Consistently, tepary bean exhibited a smaller reduction in leaf area compared to common bean under terminal drought.

Complementing the effectiveness of the vascular system, we observed that both tepary and common beans developed narrower vessels under drought. This trait has also been reported in other species subjected to water scarcity, including certain soybean genotypes, *Cedrela*, and *Syzygium aromaticum,* among others (Huang *et al*., 2022; Liu *et al.*, 2023; Nogueira *et al*., 2020; Rodríguez-Ramírez *et al*., 2022; Schreiber *et al*., 2015). As water limitation increases the risk of xylem cavitation and reduces hydraulic conductivity, this anatomical adjustment may serve as a protective strategy to limit embolism spread and maintain water transport (Cardoso *et al*., 2020; Lens *et al*., 2022). Although a reduced vessel lumen impacts fluid flow (Tombesi *et al*., 2010); this drawback may be offset by an increased number of vessels (Jang and Choi, 2018). Interestingly, we found that tepary bean increased the number of vessels in the central vein under drought. This, together with enhanced development of major veins, could compensate for the reduced vessel diameter, thereby supporting effective water distribution and transport (Nardini *et al*., 2012).

Although both species exhibited reduced stomatal conductance under drought, tepary bean increased the thickness of its leaf cuticle, in contrast to a reduction observed in common bean. While the thickening of this hydrophobic layer contributes to maintaining the plant’s water status, we cannot discard the possibility that common bean may undergo changes in wax composition, which could also effectively limit water loss (Laskós *et al*., 2021; Zhang *et al*., 2021).

### Tepary bean mitigates the impact of drought on photosynthesis though more efficient carbon assimilation

One of the earliest plant responses to water limitation is stomatal closure, which reduces water loss through transpiration but also limits CO_2_ uptake (Cornic, 2000; Liu *et al*., 2016). As expected, both species showed a decreased in stomatal conductance, under drought, resulting in reduced intercellular CO_2_ concentration (C_i_) and net photosynthesis rate (A_n_). These effects were slightly more pronounced, though not statistically significant, in common bean compared to tepary bean. Notably, under well-watered conditions, tepary bean exhibited a higher A_n_ despite lower C_i_, suggesting a more efficient CO_2_ fixation capacity than common bean. This was further supported by higher values of the maximum rate of carboxylation (V_cmax_) observed in tepary bean under the same conditions. This greater assimilation capacity was also evident in the CO_2_ response curves, where drought-stressed tepary leaves maintained a consistently higher CO_2_ assimilation rate as CO_2_ concentration increased, compared to those of common bean. Remarkably, the A_n_ values of stressed tepary bean leaves were comparable to those of non-stressed common bean, demonstrating the potential of a more effective CO_2_ assimilation in tepary bean under long drought periods. Possible explanations for these findings may involve higher Rubisco activity and/or greater Rubisco regeneration capacity, without excluding the potential contribution of increased Rubisco activase activity or other regulatory factors (Croce *et al*., 2024; Khan *et al*., 2024; Lu *et al*., 2020).

In parallel with its higher CO_2_ assimilation efficiency, we found that drought-stressed tepary bean leaves exhibited narrower palisade parenchyma cells, resulting in a higher palisade cell density compared to common bean leaves. This anatomical adjustment has been associated with a greater number of chloroplasts exposed to the intercellular air spaces, favoring CO_2_ diffusion to the sites of carbon fixation within the chloroplasts and thereby enhancing assimilation efficiency (Evans, 2021; Gago *et al*., 2020; Knauer *et al*., 2022; Olsovska *et al*., 2016).

Consistent with its higher CO_2_ assimilation rate, tepary bean accumulated greater starch levels than common bean under well-watered and drought conditions. Although both species showed a reduction in starch content under drought, the decrease was more pronounced in common bean, suggesting less abundance of the precursors for starch biosynthesis rather than higher starch degradation. Likewise, sucrose concentrations were higher in tepary bean than in common bean during drought, along with increased levels of its degradation products, glucose and fructose. This pattern suggests enhanced sucrose catabolism in tepary bean, likely supporting the development of the anatomical adjustments required for continued growth under stress and the translocation of resources to sink organs involved in reproductive processes (Kaur *et al*., 2021; Li *et al*., 2024; Liu *et al*., 2012).

When analyzing the foliage in both species, we observed better shoot biomass maintenance in *Pa* and stronger growth inhibition in *Pv* under drought, likely reflecting resource limitation, reallocation, or growth arrest to reduce transpiring leaf area (Yavas *et al.*, 2024; Zhang *et al*., 2020). Tepary bean increased its specific leaf area (SLA) in response to drought treatment, whereas common bean showed a decrease. SLA, a key functional trait describing the ratio of leaf area to dry leaf biomass, provides insight into the balance between carbon gain and water loss (Cornelissen *et al*., 2003; Liu *et al*., 2017). The contrasting responses suggest that tepary bean allocates resources to sustain biomass and maintain growth despite water scarcity. In contrast, the reduced SLA in common bean indicates stronger leaf growth inhibition, which, although limiting expansion, may help reduce transpirational surface area and promote water conservation (Kaproth *et al*., 2023; Wang *et al*., 2024). The higher SLA values observed in fully expanded Pv leaves under optimal irrigation compared to water-limited conditions also align with a lower leaf density per unit area, reflecting greater water and resource availability (Navarro and Hidalgo-Triana, 2021; Wellstein *et al*., 2017).

As a general observation, the comparative transcriptomic analysis revealed a relatively small number of DATs in *Pa* leaves and limited overlap with *Pv* DATs, consistent with previous reports indicating that *Pa* possesses drought-avoidance traits, such as greater hydraulic conductivity, deeper rooting, and smaller leaves, that support higher photosynthetic efficiency (Beebe *et al*., 2013; Rao, 2013). In contrast, *Pv* mounted a more intense stress response, evidenced by the larger number and higher accumulation of up-regulated transcripts. Notably, *Pv* showed stronger enrichment of transcripts involved in amino acid metabolism, suggesting that, under the energy deprivation imposed by the stress treatment, the leaves of this cultivar exploit amino acids not only as alternative energy substrates but also as protective molecules and key regulators of stress responses and signaling pathways (Heidarzadeh, 2025; Heinemann and Hildebrandt, 2021). *Pv* also exhibited increase abundance of transcripts related to plasma membrane turnover, which may be linked to the greater length of palisade cells observed under drought. By contrast, *Pa* transcriptomic profile indicated adjustment strategies involving cell-wall metabolism and biogenesis, as well as developmental and morphological changes likely responsible for observed leaf structural traits such as longer major veins, higher central xylem-vessel density, and a thicker cuticle.

The common bean genotype used in this study, Pinto Saltillo from Durango race, was developed for terminal drought resistance and is recognized for its exceptional yield and grain quality (Sanchez-Valdez *et al*., 2004). Previous studies show that this cultivar responds to terminal drought by efficiently translocating carbon from source leaves to pods and seeds (Cuellar-Ortiz *et al*., 2008). Physiological and molecular evidence indicates that this genotype prioritizes resource remobilization to reproductive organs over the maintenance of leaf growth and function (González-Lemes *et al*., 2023; Polania *et al.*, 2022). These findings align with our observations of reduced leaf performance in common bean under the conditions tested, compared to the tepary bean TARS-Tep32 genotype, also recognized for its resistance to terminal drought (Porch *et al*., 2013). The results presented here highlight the stronger stress response in *Pv* leaves at the reproductive stage, in correlation with its preference of sustaining pod development and functions under drought, whereas tepary bean leaves maintains leaf function through anatomical and physiological adjustments that enable continuous resource remobilization. The leaf traits identified in tepary bean provide valuable insights for enhancing drought resilience in common bean, particularly in the context of climate change.

## Supplementary data

**Supplementary Figure S1**. Measurements of leaf traits.

**Supplementary Figure S2.** Leaf relative chlorophyl content in *P. vulgaris* and *P. acutifolius* cultivars

**Supplementary Figure S3**. Fitting of A-Ci curves to the Farquhar-Berry-von Caemmerer photosynthesis model.

**Supplementary Figure S4.** Proportion of spongy parenchyma cells in leaves of *Phaseolus vulgaris* (Pv) and *Phaseolus acutifolius* (Pa).

**Supplementary Figure S5**. Comparative transcriptomic analysis of DATs of *Phaseolus vulgaris* and *Phaseolus acutifolius* leaves under terminal drought.

**Supplementary Figure S6**. Enrichment analysis of up-regulated DATs in leaf transcriptomes of *P. vulgaris* and *P. acutifolius* under well-watered conditions.

**Supplementary Figure S7**. Heat map showing the comparative abundance of shared up-regulated transcripts in response to terminal drought in both *Phaseolus* species.

**Supplementary Table S1**. List of shared transcripts in response to terminal drought according to GO classification.

**Supplementary Table S2**. List of up-regulated DATs in response to terminal drought according to KEGG pathway classification.

## Acknowledgements

We are grateful to Dr. Jorge Acosta-Gallegos and Dr. José A. Polania for their critical review of this manuscript. We thank Ing. Leticia Olvera, in Dr. Gloria Saab lab, for technical assistance with starch quantification, and Fernando González-Muñoz, in Dr. Agustín López-Munguía lab, for his generous support in the acquisition of sugar quantification data.

## Author contribution

AVS-Ch conceptualized, performed and designed the research, analyzed data and wrote the original draft; GTZ-P conducted and supervised microscopy-related experiments; AA-M supervised some of the plant growth treatments and physiological measurements; AL-J and VMP carried out the bioinformatic analyses; FG-M performed the HPLC determinations of sugar content; AAC conceptualized and supervised the research, secured funding, and wrote the manuscript with input from all authors.

## Conflict of interest

The authors declare they have no conflict of interest.

## Funding

This work was supported by the Programa de Apoyo a Proyectos de Investigación e Innovación Tecnológica (PAPIIT) de la Dirección General de Asuntos del Personal Académico (DGAPA) de la Universidad Nacional Autónoma de México (UNAM) (IN-IN209723) to AAC. This paper is part of the doctoral research conducted by VS-C, who thanks the Posgrado en Ciencias Biológicas at the Universidad Nacional Autónoma de México (UNAM) and acknowledges a scholarship from the Consejo Nacional de Ciencia y Tecnología (CONAHCyT; grant number 777020).

## Data availability

All data supporting the findings of this study are available within the paper and within its supplementary materials published online.

## Notes

### Competing Interest Statement

The authors have declared no competing interest.

## REFERENCES

AbdElgawad H, Avramova V, Baggerman G, Van Raemdonck G, Valkenborg D, Van Ostade X, Guisez Y, Prinsen E, Asard H, Van den Ende W, Beemster GTS. 2020. Starch biosynthesis contributes to the maintenance of photosynthesis and leaf growth under drought stress in maize. Plant Cell and Environment 43, 2254–2271.

Acosta-Gallegos JA, Shibata JK. 1989. Effect of Water-Stress on Growth and Yield of Indeterminate Dry-Bean (*Phaseolus vulgaris*) Cultivars. Field Crops Research 20, 81–93.

Acosta-Maspons A, Gonzalez-Lemes I, Covarrubias AA. 2019. Improved protocol for isolation of high-quality total RNA from different organs of *Phaseolus vulgaris* L. Biotechniques 66, 96–98.

Ahmad HB, Lens F, Capdeville G, Burlett R, Lamarque LJ, Delzon S. 2018. Intraspecific variation in embolism resistance and stem anatomy across four sunflower (*Helianthus annuus* L.) accessions. Physiologia Plantarum 163, 59–72.

Barrera S, Teran JC, Aparicio J, Diaz J, Leon R, Beebe S, Urrea CA, Gepts P. 2024. Identification of drought and heat tolerant tepary beans in a multi-environment trial study. Crop Science 64, 3399–3416.

Beebe SE, Rao IM, Blair MW, Acosta-Gallegos JA. 2013. Phenotyping common beans for adaptation to drought. Frontiers in Physiology 4, 35.

Beerling DJ, Franks PJ. 2010. Plant science: The hidden cost of transpiration. Nature 464, 495–496.

Bernacchi CJ, Bagley JE, Serbin SP, Ruiz-Vera UM, Rosenthal DM, Vanloocke A. 2013. Modelling C3 photosynthesis from the chloroplast to the ecosystem. Plant Cell and Environment 36, 1641–1657.

Bolger AM, Lohse M, Usadel B. 2014. Trimmomatic: a flexible trimmer for Illumina sequence data. Bioinformatics 30, 2114–2120.

Braun DM, Wang L, Ruan YL. 2014. Understanding and manipulating sucrose phloem loading, unloading, metabolism, and signalling to enhance crop yield and food security. Journal of Experimental Botany 65, 1713–1735.

Bray S, Reid DM. 2002. The effect of salinity and CO_2_ enrichment on the growth and anatomy of the second trifoliate leaf of the second trifoliate leaf of *Phaseolus vulgaris*. Canadian Journal of Botany-Revue Canadienne De Botanique 80, 349–359.

Broughton WJ, Hernández G, Blair M, Beebe S, Gepts P, Vanderleyden J. 2003. Beans (*Phaseolus* spp.) – model food legumes. Plant and Soil 252, 55–128.

Cano FJ, Sánchez-Gómez D, Rodríguez-Calcerrada J, Warren CR, Gil L, Aranda I. 2013. Effects of drought on mesophyll conductance and photosynthetic limitations at different tree canopy layers. Plant Cell and Environment 36, 1961–1980.

Cardoso AA, Batz TA, McAdam SAM. 2020. Xylem Embolism Resistance Determines Leaf Mortality during Drought in during Drought in Persea americana. Plant Physiology 182, 547–554.

Chen K. 2024. Tepary Bean (Phaseolus acutifolius). Fort Collins, Colorado: Colorado State University.

Chen MJ, Zhu XF, Zhang Y, Du ZH, Chen XB, Kong XR, Sun WJ, Chen CS. 2020. Drought stress modify cuticle of tender tea leaf and mature leaf for transpiration barrier enhancement through common and distinct modes. Scientific Reports 10.

Clarke VC, Danila FR, von Caemmerer S. 2021. CO_2_ diffusion in tobacco: a link between mesophyll conductance and leaf anatomy. Interface Focus 11.

Cornelissen JHC, Lavorel S, Garnier E, Díaz S, Buchmann N, Gurvich DE, Reich PB, ter Steege H, Morgan HD, van der Heijden MGA, Pausas JG, Poorter H. 2003. A handbook of protocols for standardised and easy measurement of plant functional traits worldwide. Australian Journal of Botany 51, 335–380.

Cornic G. 2000. Drought stress inhibits photosynthesis by decreasing stomatal aperture - not by affecting ATP synthesis. Trends in Plant Science 5, 187–188.

Croce R, Carmo-Silva E, Cho YB, Ermakova M, Harbinson J, Lawson T, McCormick AJ, Niyogi KK, Ort DR, Patel-Tupper D, Pesaresi P, Raines C, Weber APM, Zhu XG. 2024. Perspectives on improving photosynthesis to increase crop yield. Plant Cell 36, 3944–3973.

Cuellar-Ortiz SM, De La Paz Arrieta-Montiel M, Acosta-Gallegos J, Covarrubias AA. 2008. Relationship between carbohydrate partitioning and drought resistance in common bean. Plant Cell and Environment 31, 1399–1409.

de Avila Silva L, Omena-Garcia RP, Condori-Apfata JA, Costa PMA, Silva NM, DaMatta FM, Zsogon A, Araujo WL, de Toledo Picoli EA, Sulpice R, Nunes-Nesi A. 2021. Specific leaf area is modulated by nitrogen via changes in primary metabolism and parenchymal thickness in pepper. Planta 253, 16.

Debouck DG, Barrantes NC, Villalobos RA. 2024. Conservation status of three rare endemic *Phaseolus* bean (Leguminosae, Phaseoleae) species of Costa Rica. Nordic Journal of Botany 2024.

Dovrat G, Meron E, Shachak M, Golodets C, Osem Y. 2019. Plant size is related to biomass partitioning and stress resistance in water-limited annual plant communities. Journal of Arid Environments 165, 1–9.

Duursma RA. 2015. Plantecophys - An R Package for Analysing and Modelling Leaf Gas Exchange Data. Plos One 10.

Earley AM, Nolting KM, Donovan LA, Burke JM. 2024. Trait variation and performance across varying levels of drought stress in cultivated sunflower (*Helianthus annuus* L.). Aob Plants 16.

Egesa AO, Vallejos CE, Begcy K. 2024. Cell size differences affect photosynthetic capacity in a Mesoamerican and an Andean genotype of *Phaseolus vulgaris* L. Frontiers in Plant Science 15.

Evans JR. 2021. Mesophyll conductance: walls, membranes and spatial complexity. New Phytologist 229, 1864–1876.

Everard JDL, W.H. 2017. Primary Products of Photosynthesis, Sucrose and Other Soluble Carbohydrates. Encyclopedia of Applied Plant Sciences (Second Edition), Vol. 1: Elsevier, 96–104.

Ewels P, Magnusson M, Lundin S, Kaller M. 2016. MultiQC: summarize analysis results for multiple tools and samples in a single report. Bioinformatics 32, 3047–3048.

Fang Y, Xiong L. 2015. General mechanisms of drought response and their application in drought resistance improvement in plants. Cellular and Molecular Life Sciences 72, 673–689.

Farooq M, Hussain M, Siddique KHM. 2014. Drought Stress in Wheat during Flowering and Grain-filling Periods. Critical Reviews in Plant Sciences 33, 331–349.

Flexas J, Clemente-Moreno MJ, Bota J, Brodribb TJ, Gago J, Mizokami Y, Nadal M, Perera-Castro AV, Roig-Oliver M, Sugiura D, Xiong DL, Carriquí M. 2021. Cell wall thickness and composition are involved in photosynthetic limitation. Journal of Experimental Botany 72, 3971–3986.

Flexas J, Medrano H. 2002. Drought-inhibition of photosynthesis in C3 plants: Stomatal and non-stomatal limitations revisited. Annals of Botany 89, 183–189.

Funk JL, Larson JE, Ricks-Oddie J. 2021. Plant traits are differentially linked to performance in a semiarid ecosystem. Ecology 102.

Gago J, Carriquí M, Nadal M, Clemente-Moreno MJ, Coopman RE, Fernie AR, Flexas J. 2019. Photosynthesis Optimized across Land Plant Phylogeny. Trends in Plant Science 24, 947–958.

Gago J, Daloso DM, Carriquí M, Nadal M, Morales M, Araújo WL, Nunes-Nesi A, Flexas J. 2020. Mesophyll conductance: the leaf corridors for photosynthesis. Biochemical Society Transactions 48, 429–439.

Ganesan K, Xu BJ. 2017. Polyphenol-Rich Dry Common Beans (*Phaseolus vulgaris* L.) and Their Health Benefits. International Journal of Molecular Sciences 18.

Garcia-Gutierrez E, Ortega-Escalona F, Angeles G. 2020. A novel, rapid technique for clearing leaf tissues. Applications in Plant Sciences 8.

Gepts P, Aragão FJL, Barros Ed, Blair MW, Brondani R, Broughton W, Galasso I, Hernández G, Kami J, Lariguet P, McClean P, Melotto M, Miklas P, Pauls P, Pedrosa-Harand A, Porch T, Sánchez F, Sparvoli F, Yu K. 2008. Genomics of *Phaseolus* Beans, a Major Source of Dietary Protein and Micronutrients in the Tropics. In: Moore PH, Ming R, eds. Genomics of Tropical Crop Plants. New York, NY: Springer New York, 113–143.

Gimeno TE, Saavedra N, Ogée J, Medlyn BE, Wingate L. 2019. A novel optimization approach incorporating non-stomatal limitations predicts stomatal behaviour in species from six plant functional types. Journal of Experimental Botany 70, 1639–1651.

González-Lemes I, Acosta-Maspons A, Cetz-Chel JE, Polania JA, Acosta-Gallegos JA, Herrera-Estrella A, Covarrubias AA. 2023. Carbon-concentrating mechanisms in pods are key elements for terminal drought resistance in *Phaseolus vulgaris*. Journal of Experimental Botany 74, 1642–1658.

Grassi G, Magnani F. 2005. Stomatal, mesophyll conductance and biochemical limitations to photosynthesis as affected by drought and leaf ontogeny in ash and oak trees. Plant Cell and Environment 28, 834–849.

Han JM, Lei ZY, Flexas J, Zhang YJ, Carriqui M, Zhang WF, Zhang YL. 2018. Mesophyll conductance in cotton bracts: anatomically determined internal CO_2_ diffusion constraints on photosynthesis. Journal of Experimental Botany 69, 5433–5443.

Han JM, Lei ZY, Zhang YJ, Yi XP, Zhang WF, Zhang YL. 2019. Drought-introduced variability of mesophyll conductance in *Gossypium* and its relationship with leaf anatomy. Physiologia Plantarum 166, 873–887.

Haworth M, Centritto M, Giovannelli A, Marino G, Proietti N, Capitani D, De Carlo A, Loreto F. 2017. Xylem morphology determines the drought response of two *Arundo donax* ecotypes from contrasting habitats. Global Change Biology Bioenergy 9, 119–131.

Heidarzadeh A. 2025. Role of amino acids in plant growth, development, and stress responses: a comprehensive review. Discover Plants 2, 237.

Heinemann B, Hildebrandt TM. 2021. The role of amino acid metabolism in signaling and metabolic adaptation to stress-induced energy deficiency in plants. Journal of Experimental Botany 72, 4634–4645.

Himes A, Emerson P, McClung R, Renninger H, Rosenstiel T, Stanton B. 2021. Leaf traits indicative of drought resistance in hybrid poplar. Agricultural Water Management 246, 106676.

Huang WW, Fonti P, Lundqvist SO, Larsen JB, Hansen JK, Thygesen LG. 2022. Differences in xylem response to drought provide hints to future species selection. New Forests 53, 759–777.

Huang WW, Ratkowsky DA, Hui C, Wang P, Su JL, Shi PJ. 2019. Leaf Fresh Weight Versus Dry Weight: Which is Better for Describing the Scaling Relationship between Leaf Biomass and Leaf Area for Broad-Leaved Plants? Forests 10.

Jang G, Choi YD. 2018. Drought stress promotes xylem differentiation by modulating the interaction between cytokinin and jasmonic acid. Plant Signaling & Behavior 13.

Kaproth MA, Fredericksen BW, González-Rodríguez A, Hipp AL, Cavender-Bares J. 2023. Drought response strategies are coupled with leaf habit in 35 evergreen and deciduous oak (*Quercus*) species across a climatic gradient in the Americas. New Phytologist 239, 888–904.

Kaur H, Manna M, Thakur T, Gautam V, Salvi P. 2021. Imperative role of sugar signaling and transport during drought stress responses in plants. Physiologia Plantarum 171, 833–848.

Khan N, Choi SH, Lee CH, Qu MN, Jeon JS. 2024. Photosynthesis: Genetic Strategies Adopted to Gain Higher Efficiency. International Journal of Molecular Sciences 25.

Knauer J, Cuntz M, Evans JR, Niinemets Ü, Tosens T, Veromann-Jürgenson LL, Werner C, Zaehle S. 2022. Contrasting anatomical and biochemical controls on mesophyll conductance across plant functional types. New Phytologist 236, 357–368.

Krasensky J, Jonak C. 2012. Drought, salt, and temperature stress-induced metabolic rearrangements and regulatory networks. Journal of Experimental Botany 63, 1593–1608.

Kuhlgert S, Austic G, Zegarac R, Osei-Bonsu I, Hoh D, Chilvers MI, Roth MG, Bi K, TerAvest D, Weebadde P, Kramer DM. 2016. MultispeQ Beta: a tool for large-scale plant phenotyping connected to the open PhotosynQ network. R Soc Open Sci 3, 160592.

Langmead B, Salzberg SL. 2012. Fast gapped-read alignment with Bowtie 2. Nature Methods 9, 357–U354.

Laskós K, Czyczylo-Mysza IM, Dziurka M, Noga A, Góralska M, Bartyzel J, Mysków B. 2021. Correlation between leaf epicuticular wax composition and structure, physio-biochemical traits and drought resistance in glaucous and non-glaucous near-isogenic lines of rye. Plant Journal 108, 93–119.

Legner N, Fleck S, Leuschner C. 2014. Within-canopy variation in photosynthetic capacity, SLA and foliar N in temperate broad-leaved trees with contrasting shade tolerance. Trees-Structure and Function 28, 263–280.

Lei ZY, Han JM, Yi XP, Zhang WF, Zhang YL. 2018. Coordinated variation between veins and stomata in cotton and its relationship with water-use efficiency under drought stress. Photosynthetica 56, 1326–1335.

Lens F, Gleason SM, Bortolami G, Brodersen C, Delzon S, Jansen S. 2022. Functional xylem characteristics associated with drought-induced embolism in angiosperms. New Phytologist 236, 2019–2036.

Li J, He CC, Liu SH, Guo YT, Zhang YX, Zhang LJ, Zhou X, Xu DY, Luo X, Liu HY, Yang XR, Wang Y, Shi J, Yang B, Wang J, Wang PR, Deng XJ, Sun CH. 2024. Research progress and application strategies of sugar transport mechanisms in rice. Frontiers in Plant Science 15.

Li JY, Xie LF, Ren JJ, Zhang TX, Cui JH, Bao ZLT, Zhou WF, Bai J, Gong CM. 2022. CkREV regulates xylem vessel development in *Caragana korshinskii* in response to drought. Frontiers in Plant Science 13.

Liao ZG, Zhang YC, Yu Q, Fang WC, Chen MY, Li TF, Liu Y, Liu ZC, Chen L, Yu SW, Xia H, Xue HW, Yu H, Luo LJ. 2023. Coordination of growth and drought responses by GA-ABA signaling in rice. New Phytologist 240, 1149–1161.

Liu DD, Chao WM, Turgeon R. 2012. Transport of sucrose, not hexose, in the phloem. Journal of Experimental Botany 63, 4315–4320.

Liu JC, Temme AA, Cornwell WK, van Logtestijn RSP, Aerts R, Cornelissen JHC. 2016. Does plant size affect growth responses to water availability at glacial, modern and future CO_2_ concentrations? Ecological Research 31, 213–227.

Liu Q, Liu Y, Gao LQ, Wang YX, Yang MY, Wang GL. 2023. Vessel, intervessel pits and vessel-to-fiber pits have significant impact on hydraulic function under different drought conditions and re-irrigation. Environmental and Experimental Botany 214.

Liu ZL, Zhu Y, Li FR, Jin GZ. 2017. Non-destructively predicting leaf area, leaf mass and specific leaf area based on a linear mixed-effect model for broadleaf species. Ecological Indicators 78, 340–350.

Long RW, Adams HD. 2023. The osmotic balancing act: When sugars matter for more than metabolism in woody plants. Global Change Biology 29, 1684–1687.

Love MI, Huber W, Anders S. 2014. Moderated estimation of fold change and dispersion for RNA-seq data with DESeq2. Genome Biology 15.

Lu XH, Ju WM, Li J, Croft H, Chen JM, Luo YQ, Yu H, Hu HJ. 2020. Maximum Carboxylation Rate Estimation With Chlorophyll Content as a Proxy of Rubisco Content. Journal of Geophysical Research-Biogeosciences 125.

MacNeill GJ, Mehrpouyan S, Minow MAA, Patterson JA, Tetlow IJ, Emes MJ. 2017. Starch as a source, starch as a sink: the bifunctional role of starch in carbon allocation. Journal of Experimental Botany 68, 4433–4453.

Mano NA, Madore B, Mickelbart MV. 2023. Different Leaf Anatomical Responses to Water Deficit in Maize and Soybean. Life-Basel 13, 290.

Manter DK, Kerrigan J. 2004. A/Ci curve analysis across a range of woody plant species: influence of regression analysis parameters and mesophyll conductance. Journal of Experimental Botany 55, 2581–2588.

Martin M. 2011. Cutadapt removes adapter sequences from high-throughput sequencing reads. EMBnet Journal 17, 10–12.

Mendes KR, Batista-Silva W, Dias-Pereira J, Pereira MPS, Souza EV, Serrao JE, Granja JAA, Pereira EC, Gallacher DJ, Mutti PR, da Silva DTC, de Souza Junior RS, Costa GB, Bezerra BG, Silva C, Pompelli MF. 2022. Leaf plasticity across wet and dry seasons in *Croton blanchetianus* (Euphorbiaceae) at a tropical dry forest. Scientific Reports 12, 954.

Micheletto S, Rodriguez-Uribe L, Hernandez R, Richins RD, Curry J, O’Connell MA. 2007. Comparative transcript profiling in roots of *Phaseolus acutifolius* and *P. vulgaris* and under water deficit stress. Plant Science 173, 510–520.

Miller GL. 1959. Use of Dinitrosalicylic Acid Reagent for Determination of Reducing Sugar. Analytical Chemistry 31, 426–428.

Mizokami Y, Sugiura D, Watanabe CKA, Betsuyaku E, Inada N, Terashima I. 2019. Elevated CO_2_-induced changes in mesophyll conductance and anatomical traits in wild type and carbohydrate-metabolism mutants of Arabidopsis. Journal of Experimental Botany 70, 4807–4818.

Moghaddam SM, Oladzad A, Koh C, Ramsay L, Hart JP, Mamidi S, Hoopes G, Sreedasyam A, Wiersma A, Zhao D, Grimwood J, Hamilton JP, Jenkins J, Vaillancourt B, Wood JC, Schmutz J, Kagale S, Porch T, Bett KE, Buell CR, McClean PE. 2021. The tepary bean genome provides insight into evolution and domestication under heat stress. Nature Communications 12, 2638.

Money NP. 1989. Osmotic Pressure of Aqueous Polyethylene Glycols : Relationship between Molecular Weight and Vapor Pressure Deficit. Plant Physiology 91, 766–769.

Moreno A, Damian-Almazo JY, Miranda A, Saab-Rincon G, Gonzalez F, Lopez-Munguia A. 2010. Transglycosylation reactions of α-amylase. Enzyme and Microbial Technology 46, 331–337.

Nadal M, Flexas J. 2018. Chapter 17 - Mesophyll Conductance to CO_2_ Diffusion: Effects of Drought and Opportunities for Improvement. In: García Tejero IF, Durán Zuazo VH, eds. Water Scarcity and Sustainable Agriculture in Semiarid Environment: Academic Press, 403–438.

Nardini A, Pedà G, La Rocca N. 2012. Trade-offs between leaf hydraulic capacity and drought vulnerability: morpho-anatomical bases, carbon costs and ecological consequences. New Phytologist 196, 788–798.

Navarro T, Hidalgo-Triana N. 2021. Variations in Leaf Traits Modulate Plant Vegetative and Reproductive Phenological Sequencing Across Arid Mediterranean Shrublands. Frontiers in Plant Science 12.

Niinemets Ü, Díaz-Espejo A, Flexas J, Galmés J, Warren CR. 2009. Importance of mesophyll diffusion conductance in estimation of plant photosynthesis in the field. Journal of Experimental Botany 60, 2271–2282.

Nogueira M, Livingston D, Tuong T, Sinclair TR. 2020. Xylem vessel radii comparison between soybean genotypes differing in tolerance to drought. Journal of Crop Improvement 34, 404–413.

Olsovska K, Kovar M, Brestic M, Zivcak M, Slamka P, Shao HB. 2016. Genotypically Identifying Wheat Mesophyll Conductance Regulation under Progressive Drought Stress. Frontiers in Plant Science 7.

Ozturk M, Unal BT, García-Caparrós P, Khursheed A, Gul A, Hasanuzzaman M. 2021. Osmoregulation and its actions during the drought stress in plants. Physiologia Plantarum 172, 1321–1335.

Polania JA, Salazar-Chavarria V, Gonzalez-Lemes I, Acosta-Maspons A, Chater CCC, Covarrubias AA. 2022. Contrasting *Phaseolus* Crop Water Use Patterns and Stomatal Dynamics in Response to Terminal Drought. Frontiers in Plant Science 13.

Porch TG, Beaver JS, Brick MA. 2013. Registration of Tepary Germplasm with Multiple-Stress Tolerance, TARS-Tep 22 and TARS-Tep 32. Journal of Plant Registrations 7, 358–364.

Porch TG, Cichy K, Wang WJ, Brick M, Beaver JS, Santana-Morant D, Grusak MA. 2017. Nutritional composition and cooking characteristics of tepary bean (*Phaseolus acutifolius* Gray) in comparison with common bean (*Phaseolus vulgaris* L.). Genetic Resources and Crop Evolution 64, 935–953.

Rao IM, Beebe, S. E., Polania, J., Ricaute, J., Cajiao, C., Garcia, R., Rivera, M. 2013. Can tepary bean be a model for improvement of drought resistance in common bean? African Crop Science Journal 21, 265–281.

Rodríguez-Ramírez EC, Ferrero ME, Acevedo-Vega I, Crispin-DelaCruz DB, Ticse-Otarola G, Requena-Rojas EJ. 2022. Plastic adjustments in xylem vessel traits to drought events in three *Cedrela* species from Peruvian Tropical Andean forests. Scientific Reports 12.

Romero MF, Gallego D, Lechuga-Jiménez A, Martínez JF, Barajas HR, Hayano-Kanashiro C, Peimbert M, Cruz-Ortega R, Molina-Freaner FE, Alcaraz LD. 2021. Metagenomics of mine tailing rhizospheric communities and its selection for plant establishment towards bioremediation. Microbiological Research 247.

Rosales MA, Cuellar-Ortiz SM, de la Paz Arrieta-Montiel M, Acosta-Gallegos J, Covarrubias AA. 2013. Physiological traits related to terminal drought resistance in common bean (*Phaseolus vulgaris* L.). Journal of the Science of Food and Agriculture 93, 324–331.

Ruan YL, Jin Y, Yang YJ, Li GJ, Boyer JS. 2010. Sugar Input, Metabolism, and Signaling Mediated by Invertase: Roles in Development, Yield Potential, and Response to Drought and Heat. Molecular Plant 3, 942–955.

Sack L, Scoffoni C. 2013. Leaf venation: structure, function, development, evolution, ecology and applications in the past, present and future. New Phytologist 198, 983–1000.

Saeed N, Maqbool N, Haseeb M, Sadiq R. 2016. Morpho-anatomical changes in roots of chickpea (*Cicer arietinum* L.) under drought stress condition. Journal of Agricultural Science and Technology 6, 1–9.

Sanchez-Valdez I, Acosta-Gallegos JA, Ibarra-Perez FJ, Rosales-Serna R, Singh SP. 2004. Registration of ‘Pinto Saltillo’ common bean. Crop Science 44, 1865+.

Schreiber SG, Hacke UG, Hamann A. 2015. Variation of xylem vessel diameters across a climate gradient: insight from a reciprocal transplant experiment with a widespread boreal tree. Functional Ecology 29, 1392–1401.

Scoffoni C, Rawls M, McKown A, Cochard H, Sack L. 2011. Decline of Leaf Hydraulic Conductance with Dehydration: Relationship to Leaf Size and Venation Architecture. Plant Physiology 156, 832–843.

Shafqat W, Mazrou YSA, Nehela Y, Ikram S, Bibi S, Naqvi SA, Hameed M, Jaskani MJ. 2021. Effect of Three Water Regimes on the Physiological and Anatomical Structure of Stem and Leaves of Different *Citrus* Rootstocks with Distinct Degrees of Tolerance to Drought Stress. Horticulturae 7.

Shao YJ, Lin AHM. 2018. Improvement in the quantification of reducing sugars by miniaturizing the Somogyi-Nelson assay using a microtiter plate. Food Chemistry 240, 898–903.

Smith MR, Veneklaas E, Polania J, Rao IM, Beebe SE, Merchant A. 2019. Field drought conditions impact yield but not nutritional quality of the seed in common bean (*Phaseolus vulgaris* L.). Plos One 14, e0217099.

Song XY, Zhou GS, He QJ, Zhou HL. 2021. Quantitative Response of Maize Vcmax25 to Persistent Drought Stress at Different Growth Stages. Water 13.

Suarez JC, Contreras AT, Anzola JA, Vanegas JI, Rao IM. 2022. Physiological Characteristics of Cultivated Tepary Bean (*Phaseolus acutifolius* A. Gray) and Its Wild Relatives Grown at High Temperature and Acid Soil Stress Conditions in the Amazon Region of Colombia. Plants-Basel 11.

Thomas PD, Kejariwal A, Campbell MJ, Mi H, Diemer K, Guo N, Ladunga I, Ulitsky-Lazareva B, Muruganujan A, Rabkin S, Vandergriff JA, Doremieux O. 2003. PANTHER: a browsable database of gene products organized by biological function, using curated protein family and subfamily classification Nucleic Acids Research 31, 334.

Tombesi S, Johnson RS, Day KR, DeJong TM. 2010. Relationships between xylem vessel characteristics, calculated axial hydraulic conductance and size-controlling capacity of peach rootstocks. Annals of Botany 105, 327–331.

van Daalen KR, Romanello M, Rocklöv J. 2022. The 2022 Europe report of the Lancet Countdown on health and climate change: towards a climate resilient future (vol 7, pg e942, 2022). Lancet Public Health 7, E993–E993.

Varone L, Ribas-Carbo M, Cardona C, Gallé A, Medrano H, Gratani L, Flexas J. 2012. Stomatal and non-stomatal limitations to photosynthesis in seedlings and saplings of Mediterranean species pre-conditioned and aged in nurseries: Different response to water stress. Environmental and Experimental Botany 75, 235–247.

Velikova V, Arena C, Izzo LG, Tsonev T, Koleva D, Tattini M, Roeva O, De Maio A, Loreto F. 2020. Functional and Structural Leaf Plasticity Determine Photosynthetic Performances during Drought Stress and Recovery in Two *Platanus orientalis* Populations from Contrasting Habitats. International Journal of Molecular Sciences 21.

Vicente-Serrano SM, Peña-Angulo D, Beguería S, Domínguez-Castro F, Tomás-Burguera M, Noguera I, Gimeno-Sotelo L, El Kenawy A. 2022. Global drought trends and future projections. Philosophical Transactions of the Royal Society a-Mathematical Physical and Engineering Sciences 380.

Vital RG, Müller C, Freire FBS, Silva FB, Batista PF, Fuentes D, Rodrigues AA, Moura LMF, Daloso DM, Silva AA, Merchant A, Costa AC. 2022. Metabolic, physiological and anatomical responses of soybean plants under water deficit and high temperature condition. Scientific Reports 12.

Wang S, Zhou H, He ZB, Ma DK, Sun WH, Xu XZ, Tian QY. 2024. Effects of Drought Stress on Leaf Functional Traits and Biomass Characteristics of *Atriplex canescens*. Plants-Basel 13.

Weatherley PE. 1950. Studies in the Water Relations of the Cotton Plant. I. The Field Measurement of Water Deficits in Leaves. The New Phytologist 49, 81–97.

Wellstein C, Poschlod P, Gohlke A, Chelli S, Campetella G, Rosbakh S, Canullo R, Kreyling J, Jentsch A, Beierkuhnlein C. 2017. Effects of extreme drought on specific leaf area of grassland species: A meta-analysis of experimental studies in temperate and sub-Mediterranean systems. Global Change Biology 23, 2473–2481.

Yavas I, Jamal MA, Ul Din K, Ali S, Hussain S, Farooq M. 2024. Drought-Induced Changes in Leaf Morphology and Anatomy: Overview, Implications and Perspectives. Polish Journal of Environmental Studies 33, 1517–1530.

Yeats TH, Rose JKC. 2013. The Formation and Function of Plant Cuticles. Plant Physiology 163, 5–20.

Yuan X, Wang YM, Ji P, Wu PL, Sheffield J, Otkin JA. 2023. A global transition to flash droughts under climate change. Science 380, 187–191.

Zhang H, Zhao Y, Zhu JK. 2020. Thriving under Stress: How Plants Balance Growth and the Stress Response. Developmental Cell 55, 529–543.

Zhang Y, Du ZH, Han YT, Chen XB, Kong XR, Sun WJ, Chen CS, Chen MJ. 2021. Plasticity of the Cuticular Transpiration Barrier in Response to Water Shortage and Resupply in *Camellia sinensis*: A Role of Cuticular Waxes. Frontiers in Plant Science 11.

